# The E3 ubiquitin ligase RNF115 regulates phagosome maturation and host response to bacterial infection

**DOI:** 10.1101/2021.07.13.452284

**Authors:** Orsolya Bilkei-Gorzo, Tiaan Heunis, José Luis Marín-Rubio, Francesca Romana Cianfanelli, Benjamin Bernard Armando Raymond, Joseph Inns, Daniela Fabrikova, Julien Peltier, Fiona Oakley, Ralf Schmid, Anetta Härtlova, Matthias Trost

## Abstract

Phagocytosis is a key process in innate immunity and homeostasis. After uptake, newly formed phagosomes mature by acquisition of endo-lysosomal enzymes. Macrophage activation by interferon-gamma (IFN-γ) increases microbicidal activity, but delays phagosomal maturation by an unknown mechanism. Using quantitative proteomics, we show that phagosomal proteins harbour high levels of typical and atypical ubiquitin chain types. Moreover, phagosomal ubiquitylation of vesicle trafficking proteins is substantially enhanced upon IFN-γ activation of macrophages, suggesting a role in regulating phagosomal functions. We identified the E3 ubiquitin ligase RNF115, which is enriched on phagosomes of IFN-γ activated macrophages, as an important regulator of phagosomal maturation. Loss of RNF115 protein or ligase activity enhanced phagosomal maturation and increased cytokine responses to bacterial infection, suggesting that both innate immune signalling from the phagosome and phagolysosomal trafficking are controlled through ubiquitylation. RNF115 knock-out mice show less tissue damage in response to *S. aureus* infection, indicating a role of RNF115 in inflammatory responses *in vivo*. In conclusion, RNF115 and phagosomal ubiquitylation are important regulators of innate immune functions during bacterial infections.

## Introduction

Phagocytosis is an essential component of the innate immune response against invading pathogens and tissue injury (Brown et al., 2015, Pauwels et al., 2017). It is an evolutionarily conserved process by which microbes are internalised and delivered to the phagosome (Boulais et al., 2010). After internalization, the newly formed phagosome is constantly remodelled by fusion and fission processes with early and late endosomes and finally with the lysosome (Kinchen & Ravichandran, 2008). These ultimate changes deliver the engulfed pathogen into the terminal degradative compartments known as phagolysosomes. This process is highly controlled, and the activation status of the macrophage changes the process significantly (Guo et al., 2019, Pauwels et al., 2019, Trost et al., 2009). As dysregulation of phagosome maturation can lead to infectious or inflammatory disease (Jain et al., 2019), it is of great of importance to understand how phagocytosis and phagosome maturation are controlled.

Ubiquitylation is an important post-translational modification that involves the covalent attachment of ubiquitin to (mostly) lysine (K) residues of target proteins (Heap et al., 2017, Yau & Rape, 2016). This process is catalysed by a three-step enzymatic cascade comprising the E1 activating enzyme, E2 ubiquitin-conjugating enzyme and E3 ubiquitin-ligating enzymes which can be reversed by deubiquitylases (DUBs) (Komander & Rape, 2012, Ritorto et al., 2014). Ubiquitin itself can be ubiquitylated at one of its seven lysine residues and the N-terminal methionine (K6, K11, K27, K29, K33, K48, K63 and M1), leading to the assembly of polyubiquitin chains (Kulathu & Komander, 2012). These chains lead to different biological outcomes: e.g., K48 chains are associated with proteasomal degradation, whilst K63 chains are known to be involved in cell signalling and trafficking (Erpapazoglou et al., 2014, Guo et al., 2019). Ubiquitin signalling is integral to almost all cellular processes in eukaryotes and thus, disorders or mutations in ubiquitin pathways result in a wide range of diseases (Rape, 2018).

Previous work has shown that polyubiquitin chains associate with phagosomes (Lee et al., 2005). Moreover, several intracellular pathogens such as *Legionella pneumophila*, *Shigella flexneri* or *Salmonella* secrete ubiquitylation modifying enzymes (Ashida et al., 2014, Maculins et al., 2016, Qiu & Luo, 2017), suggesting that ubiquitylation regulates key processes in phagosome biology and innate immunity (Dean et al., 2019). Moreover, our proteomics data has shown that ubiquitin was enriched on phagosomes in response to IFN-γ activation (Naujoks et al., 2016, Trost et al., 2009), therefore we hypothesised that ubiquitylation may play a role in regulating phagosome functions.

Interferon-γ (IFN-γ) is essential against for host defence against intracellular infection by activating macrophages to kill bacteria as well as up-regulation of antigen processing and presentation pathways (Naujoks et al., 2016, Trost et al., 2009). Recently, we and others have shown that IFN-γ activation of macrophages delays phagosomal maturation but increases phagocytic update of particles (Trost et al., 2009, Yates et al., 2007). Although it has been shown that the IFN-induced increase of reactive oxygen species (ROS) generated by the NADPH oxidase (NOX2) complex inhibits phagosomal proteolysis (Rybicka et al., 2010, Savina et al., 2006), the exact mechanism how IFN-γ activation affects phagosomal maturation, is unknown.

Here, we show that phagosomal proteins are ubiquitylated and IFN-γ activation of macrophages substantially increases this further. We identified the E3 ligase Ring Finger Protein 115 (RNF115), which increasingly locates to phagosomes upon IFN-γ activation, as a major regulator of phagosomal functions. Loss of RNF115 affected several phagosomal vesicle trafficking pathways and increased phagosomal maturation. Moreover, loss of RNF115 also increased cytokine responses and affected infection-induced tissue damage *in vivo*. These results suggest a key role for RNF115 in phagosome functions and inflammatory responses.

## Results

### Ubiquitylation is abundant on phagosomes and increased by IFN-γ activation

As shown before, IFN-γ activation of macrophages delays phagosomal maturation but increases phagocytic update of particles (Trost et al., 2009, Yates et al., 2007) **(Figure 1A).** To investigate the role of ubiquitylation in regulating phagosome maturation, we examined the presence of polyubiquitylation in total cell lysates (TCL) and on purified phagosomes (Hartlova et al., 2017) **(Figure 1B)** in the murine macrophage cell line RAW264.7 (Guo et al., 2015). Immunoblots revealed significant enrichment of polyubiquitylated protein chains on phagosomes compared to TCL **(Figure 1C).** Moreover, inflammatory macrophage activation by IFN-γ substantially enhanced ubiquitylation of phagosomal proteins. Ubiquitylation in the endo-lysosomal system is often considered to be primarily important for protein degradation (Haglund & Dikic, 2012). However, total ubiquitylation stays almost constant over the whole phagosomal maturation process of both resting and IFN-γ-treated macrophages **(Figure 1D),** suggesting that polyubiquitin chains may be heavily involved in endo-lysosomal processes themselves and may have other roles beside degradation, i.e. serving as signalling platforms (Guo et al., 2019).

**Figure 1:**
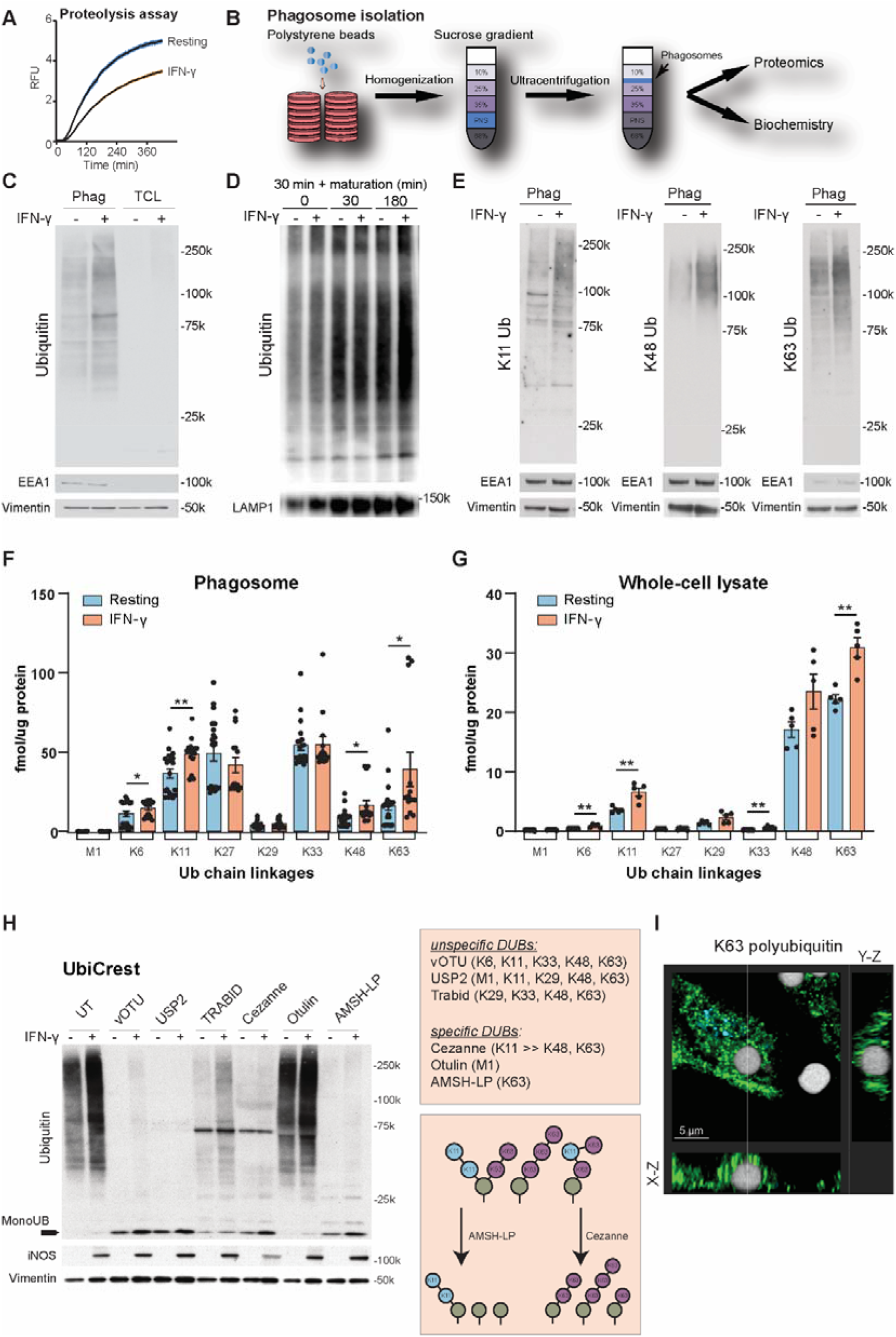
Characterisation of phagosomal ubiquitylation. (A) Intraphagosomal proteolysis assay shows that IFN-γ activation of macrophages reduces phagosomal maturation. Shaded area represents SEM. (B) Workflow of phagosome isolation. (C-E) Western blots showing increased polyubiquitylation on phagosomes compared to Total Cell Lysate (TCL) and further increases by IFN-γ activation (C, E). Equal protein amounts were loaded for phagosomal and TCL samples. Vimentin serves as loading control; EEA1 as purity marker. (D) Phagosomal ubiquitylation is not changing substantially over the maturation of the phagosome (30 min pulse, 0, 30, and 180 min chase, respectively). LAMP1 serves as purity marker. Representative images of three replicates. (F) Ubiquitin AQUA PRM assay for phagosomes from RAW264.7 cells shows abundance of atypical chains and increases for K6, K11, K48 and K63 chains on phagosomes in response to IFN-γ activation. As the experiment is complex and has inherent variability, five independent experiments of three biological replicates were combined. M1 represents linear ubiquitin chains. M1= linear ubiquitin chains. (G) Ubiquitin AQUA PRM assay of total cell lysates of RAW264.7 cells. Error bars represent SEM. *=p<0.05; **=p<0.01 by paired two-tailed Student’s t-test. (H) Ubiquitin Chain Restriction (UbiCRest) experiment removing polyubiquitin chains from phagosomal extracts using specific and unspecific DUBs. UT = untreated (I) Representative immunofluorescence micrograph showing that K63 polyubiquitin localises around the phagosome. Bead size: 3 μm.

To better understand the role of ubiquitylation of phagosomal proteins, we first determined which ubiquitin chain types were enriched on phagosomes. Comparative analysis using ubiquitin-chain type specific antibodies revealed that IFN-γ significantly induced K11, K48 and K63 polyubiquitylation chains on phagosomes **(Figure 1E).** As there are no selective antibodies for atypical ubiquitin chain types, we additionally used a quantitative targeted mass spectrometry approach, the AQUA ubiquitin Parallel Reaction Monitoring (PRM) assay **(Figure 1F/G).** This highly sensitive method enables the detection and quantification of endogenous ubiquitin chains (Heunis et al., 2020, Tsuchiya et al., 2013). The data confirmed increases of K11, K48 and K63 chains and additionally showed substantial amounts of non-canonical ubiquitin chains such as K27 and K33, whose biological function is less well understood (Kulathu & Komander, 2012, van Huizen & Kikkert, 2019) **(Figure 1F).** In contrast, K63, K48 and K11 chains dominate ubiquitin chains of TCL **(Figure 1G).** To further validate the PRM data, we performed deubiquitylase (DUB)-based analyses of phagosomal ubiquitin chain composition using Ubiquitin Chain Restriction (UbiCRest) (Hospenthal et al., 2015). In these experiments, phagosomal extracts were treated with ubiquitin-chain specific DUBs such as USP2 (unspecific), vOTU (unspecific), Otulin (specificity: M1/linear), Cezanne (specificity: K11>>K63), Trabid (specificity: K29, K33>K63) or AMSH-LP (specificity: K63) (Kristariyanto et al., 2015, Ritorto et al., 2014) and examined for the amount of monoubiquitin generated **(Figure 1H).**

Consistent with the PRM data, UbiCrest analysis revealed that the K11, K48 and K63-specific DUB Cezanne hydrolysed most of polyubiquitin chains on phagosomes. Moreover, the K63-specific DUB AMSH-LP was the most effective in hydrolysing polyubiquitin chains on phagosomes. These data indicate that K63 polyubiquitylation could form the backbone of polyubiquitin chains on phagosome from which other ubiquitin chain-types branch off. Furthermore, K63 polyubiquitylation is detectable by immunofluorescence microscopy around the phagosome of resting macrophages **(Figure 1I).**

K63-linked polyubiquitylation of proteins has been described to lead to lysosomal degradation and contributes to signal transduction and protein-protein interactions (Wu & Karin, 2015). Next, we tested whether polyubiquitin chains localize at the extra-luminal/cytoplasmic surface of phagosomes as opposed to being sequestered within phagosomes. Intact phagosomes were treated with different DUBs to hydrolyse ubiquitin chains on the surface **(Supplementary Figure 1A).** This assay is based on the notion that ubiquitylated proteins within the phagosomal lumen resist ubiquitin chain hydrolysis when subjected to controlled DUB exposure. Exposure of intact phagosomes to the unspecific DUB USP2 showed monoubiquitin in the supernatant but not in the lysate, indicating that polyubiquitin chains are mainly localized on the cytoplasmic surface of phagosomes **(Supplementary Figure 1B-C).** IFN-γ treatment increased the amount of hydrolysed polyubiquitin chains in the supernatants. The presence of vimentin, an abundant cytoskeletal protein associated with the phagosomal membrane, demonstrated that phagosomes remained intact during DUB treatment. These data demonstrate that polyubiquitin chains are mainly localized on the extra-luminal surface of phagosomes, thereby allowing an interaction with other signalling proteins.

Altogether, these data indicate that polyubiquitylation is abundant on phagosomes of resting macrophages and significantly enhanced by IFN-γ activation.

### Phagosome proteomics identifies increased ubiquitylation of vesicle trafficking proteins in response to IFN-γ

To further identify components of the ubiquitin system involved in the regulation of phagosome function in IFN-γ activated macrophages, we analysed the phagosomal proteome of resting and IFN-γ activated macrophages by a quantitative mass spectrometry approach (Breyer et al., 2021, Dill et al., 2015, Guo et al., 2019, Hartlova et al., 2018) **(Figure 2A, Supplementary Table 1)**. We identified several proteins of the ubiquitin system, including ubiquitin and ubiquitin-like modifiers, DUBs, E1 and E2 enzymes **(Figure 2B)** as well as E3 ligases **(Figure 2C).** We observed increased abundance of E1 and E2 enzymes, as well as E3 ligases, suggesting that the increased polyubiquitylation is a result of increased enzyme abundance rather than reduced deubiquitylation; particularly, since DUB abundance was overall not decreased.

**Figure 2:**
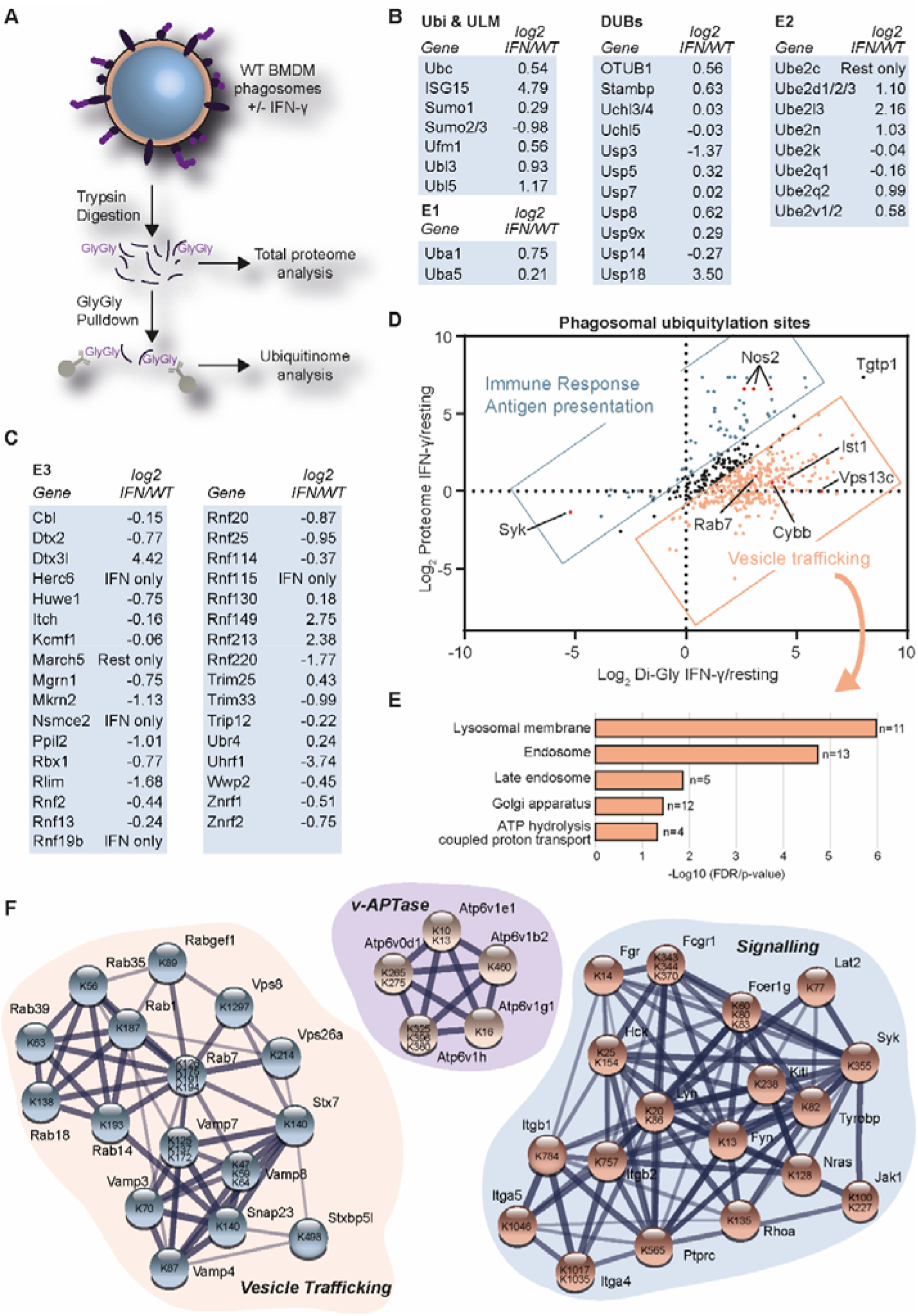
Proteomics shows that IFN-γ activation increases ubiquitylation of innate immunity and vesicle trafficking proteins. (A) Workflow. 30 min old phagosomes were isolated from RAW264.7 cells. Phagosome extracts were tryptically digested for total proteome analysis or Gly-Gly pulldowns were performed to enrich for ubiquitylated peptides. (B & C) Selected phagosomal proteins of the ubiquitin system (Ubiquitin & ubiquitin-like modifiers (ULM), E1, E2 and E3 and deubiquitylases (DUBs)) and their abundance changes in response IFN-γ activation. “IFN only” or “Rest(ing) only” are used when the proteins were only detected in the respective sample by label-free proteomics. (D) Combined analysis of total proteome and Gly-Gly proteomics data sets of phagosomes from untreated and IFN-γ activated RAW264.7 cells. (E) Gene Ontology (GO) Analysis of proteins that were ubiquitylated in response to IFN-γ activation but did not change (−0.5>Log2<0.5) significantly on the protein level. (F) Selected networks of proteins and their ubiquitylation sites obtained from STRING using data in (D). Level of shading of connecting lines indicates interaction confidence.

After tryptic digestion, ubiquitylation leaves a di-glycine (Gly-Gly) tag on the ε-amino group of lysines. Commercial antibodies allow the enrichment of these peptides for mass spectrometric analysis. We performed such a Gly-Gly pulldown of ubiquitylated peptides from 300 μg of phagosomal extracts (^~^200 cell culture dishes) per replicate. Quantitative mass spectrometry revealed 478 ubiquitylation sites on phagosomal proteins **(Supplementary Table 2).** Comparing the abundance of these Gly-Gly-modified peptides with the protein abundance changes on the total protein level showed that most proteins were overall more ubiquitylated in response to IFN-γ. Proteins that were increased in protein abundance and protein ubiquitylation were mostly involved in inflammatory and interferon-regulated responses, e.g., the nitric oxide synthase NOS2. However, interestingly, many proteins were not regulated by protein abundance but increased in ubiquitylation suggesting that they are specifically regulated by ubiquitylation. These included the endo-lysosomal Rab7, the SNARE proteins VAMP8 and Syntaxin 7 (STX7), and the NADPH oxidase 2 subunit Cybb **(Figure 2D).** Gene Ontology (GO) analysis revealed that these proteins were mostly involved in vesicle trafficking on lysosomal, endosomal and Golgi membranes as well as members of the v-ATPase complex **(Figure 2E).**

**Figure 2F** highlights networks of phagosomal proteins involved in vesicle trafficking and signalling as well as the vATPase complex that were found to be ubiquitylated. While some of the ubiquitylation sites were conserved within the sequence of substrates, for example the sites on the cytosolic tails of Integrins Itgb1 and Itgb2, others appeared to be spread over the whole sequence. Interestingly, ubiquitylation of many of the vesicle trafficking proteins (such as the VAMP proteins) appears in functional domains, suggesting that these modifications may affect the binding of their SNARE domains **(Supplementary Figure 2A).** Moreover, we realised that ubiquitylation affects recognition of antibodies in Western Blot experiments. Rab7, for example, a master regulator of endo-lysosomal trafficking (Bucci et al., 2000), is highly ubiquitylated at the C-terminal part of the protein. As this is the area against which our antibodies were raised, they do not recognise the ubiquitylated form. So Rab7 is only detectable by Western Blot in Tab2-TUBE pulldowns following removal of ubiquitylation by recombinant DUBs **(Supplementary Figure 2B).** Altogether, this dataset provides an important resource of ubiquitylation targets in the endo-lysosomal system.

### E3 ligase RNF115 locates to the phagosome and loss of RNF115 affects vesicle trafficking to the phagosome

Next, we hypothesised that the increased ubiquitylation on the phagosome was mediated by specific IFN-γ activated E3 ligases. We identified five significantly enriched E3 ligases on the phagosome in response to IFN-γ stimulation, including the interferon-induced ubiquitin-like modifier ISG15 E3 ligase Herc6 **(Figure 3A, Figure 2C).** The enriched E3 ligases included Dtx3l, which forms a complex with PARP9 and plays a role in DNA damage and antiviral responses (Zhang et al., 2015); RNF149, an uncharacterised RING ligase; RNF213, a 591 kDa multi-domain protein which has been implicated in angiogenesis, Moyamoya disease, the sensing of ISGylated proteins (Thery et al., 2021), and the ubiquitylation of LPS of intracellular bacteria (Otten et al., 2021), and the RING E3 ligase RNF115 (also called BCA2 or Rabring7) **(Figure 3A).**

**Figure 3:**
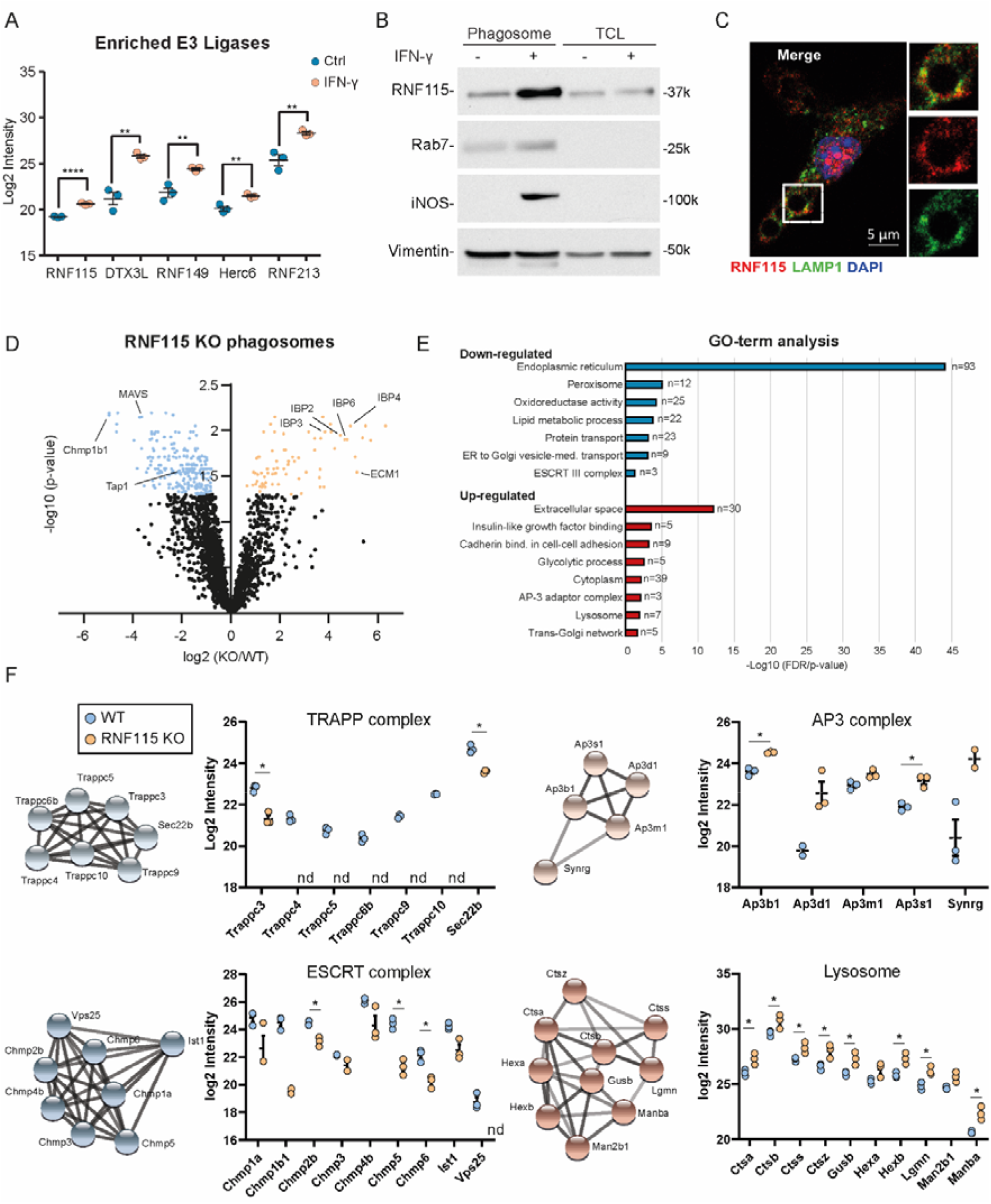
Ubiquitin E3 ligase RNF115 is enriched on phagosomes of IFN-γ activated macrophages and loss of RNF115 affects several phagosomal functions. (A) Proteomics intensity levels of selected phagosomal E3 ligases enriched upon IFN-γ activation. Missing values for RNF115 and Herc6 in resting macrophages have been added by imputation. (B) Western blot of phagosomal and TCL extracts showing enrichment of RNF115 on phagosomes of RAW264.7 cells and its increased abundance in response to IFN-γ activation. Rab7 serves as purity control, iNOS as activation control and vimentin as loading control. Representative image of three replicates. (C) Representative immunofluorescence micrograph showing RNF115 co-localising with LAMP1 around the phagosome in murine BMDMs. Scale bar equals 5 microns. (D) Volcano plot of proteomics data of resting WT and RNF115 KO BMDM phagosomes. Selected proteins are highlighted. (E) Gene Ontology (GO) analysis of enriched and depleted proteins from (D). (F) Proteomics intensity data of selected protein complexes derived from (D). Error bars represent SEM. nd = not detected. *=p<0.05.

As RNF115 was previously reported to bind to Rab7, and play a role in receptor trafficking and anti-viral host response by interfering with endo-lysosomal pathways (Li et al., 2020, Miyakawa et al., 2009, Nityanandam & Serra-Moreno, 2014, Zhang et al., 2020), we tested if RNF115 regulated phagosomal functions.

To analyse the association of RNF115 with phagosomes, we conducted immunoblot analysis of phagosomal and TCL fractions, which confirmed enrichment of RNF115 on phagosomes of IFN-γ activated and resting macrophages compared to corresponding TCLs of resting macrophages **(Figure 3B).** Consistent with immunoblot data, immunofluorescence microscopy analysis revealed RNF115 translocation on phagosomes by partly co-localising with lysosomal-associated membrane protein 1, LAMP1, in IFN-γ activated macrophages **(Figure 3C).**

To further distinguish whether RNF115 is present on the extra-luminal surface of phagosomes or if it is an intraluminal cargo, we treated isolated phagosomes with increasing amounts of trypsin to digest phagosome-associated proteins in a dose-dependent manner **(Supplementary Figure 3A).** Exposure to 0.5 μg trypsin led to a significant reduction in RNF115, similar to phagosome-associated Rab5, whereas the internal membrane protein LAMP1 resisted trypsin treatment. These data indicate that RNF115 localizes to the extraluminal surface of phagosomes.

To characterise the role of RNF115 on the phagosome, we generated a RNF115 knock-out (KO) cell line in BMA3.1A7 macrophages using CRISPR/Cas9 genome editing. The efficiency of KO was confirmed by sequencing and immunoblot analyses **(Supplementary Figure 3B).** To determine whether RNF115 is involved in the regulation of phagocytosis, we measured the uptake of green fluorescent carboxylated particles in WT and RNF115 KO macrophages. We complemented the KO cell line with HA-tagged wildtype RNF115-HA or a RNF115-HA (W259A) mutant that was predicted to be unable to bind to E2 enzymes (Hodson et al., 2014). Neither ablation of RNF115, nor expression of the mutant RNF115, affected phagocytic uptake of either carboxylated particles **(Supplementary Figure 3C)** or bacteria **(Supplementary Figure 3D).**

In order to achieve a systems level understanding of the effects of RNF115 loss, we isolated phagosomes from WT and RNF115 KO macrophages and performed proteomics analysis **(Figure 3D, Supplementary Table 3).** The phagosome, as part of the endo-lysosomal system, interacts with practically all vesicle trafficking pathways of the cell and therefore allows identifying specific pathways that RNF115 may play a role in. Loss of RNF115 affected several vesicle trafficking pathways including an increase of lysosomal and secretory proteins in the phagosome and a reduction of proteins of the ER and the peroxisome **(Figure 3E).** Of the secreted proteins, the Insulin-like growth factor-binding proteins (IGFBPs), appeared particularly retained in the endo-lysosomal system upon RNF115 loss, suggesting a role of RNF115 in correctly trafficking these proteins. Moreover, loss of RNF115 also decreased recruitment of the ESCRT complex, highlighting a potential functional role for this complex in ubiquitylation.

Other protein complexes affected by loss of RNF115 were the Transport Protein Particle (TRAPP) complex, which is involved in Rab GTPase activation (Riedel et al., 2018) and ER-Golgi and Golgi-plasma membrane trafficking (i.e. secretion) (Kim et al., 2016). TRAPP was significantly less abundant on RNF115 KO phagosomes, which may explain the retention of cargo with secretion signal **(Figure 3F)** (Zappa et al., 2019). On the other hand, the AP-3 complex, that shuttles proteins to the lysosome, is enriched in RNF115 KO phagosomes. Finally, we identified a significant increased abundance of lysosomal proteins in the RNF115 KO phagosome, suggesting that loss of RNF115 also increases phagosomal maturation **(Figure 3F).**

Altogether, this data demonstrates that loss of RNF115 affects ER-Golgi, secretory and lysosomal vesicle trafficking pathways to the phagosome and that ubiquitylation plays an important role in protein and vesicle trafficking.

### Loss of RNF115 increases phagosomal maturation

As the loss of RNF115 affected abundance of lysosomal proteins on the phagosome, we next investigated whether RNF115 was involved in the regulation of phagosomal maturation. To test the effect on phagosomal maturation, we measured phagosomal proteolytic activity and pH in WT and RNF115 KO cells using real-time quantitative fluorescent DQ-red BSA-coated particles that fluoresce at high proteolytic activity and beads coated with pHrodo dyes that fluoresce brightly in an acidic environment **(Figure 4A-B).** Ablation of RNF115 enhanced both phagosomal proteolysis and acidification, indicating that RNF115 is a negative regulator of phagosome maturation. Bafilomycin, a vacuolar ATPase inhibitor was used as a negative control for acidification measurements. IFN-γ activation also showed decreased phagosome maturation in RNF115 KO cells but not to the extent of WT cells **(Figure 4A).** This indicates that proteins other than RNF115 also play a role in regulating IFN-γ-induced changes to phagosome maturation. The importance of RNF115 in these processes was further validated by complementing RNF115 KO cells with WT and the W259A RNF115 mutant. This “ligase dead” mutant increased proteolytic activity in macrophages, similar to the KO cells, demonstrating that RNF115 E3 ligase activity is needed for regulation of phagosome maturation **(Figure 4C).**

**Figure 4:**
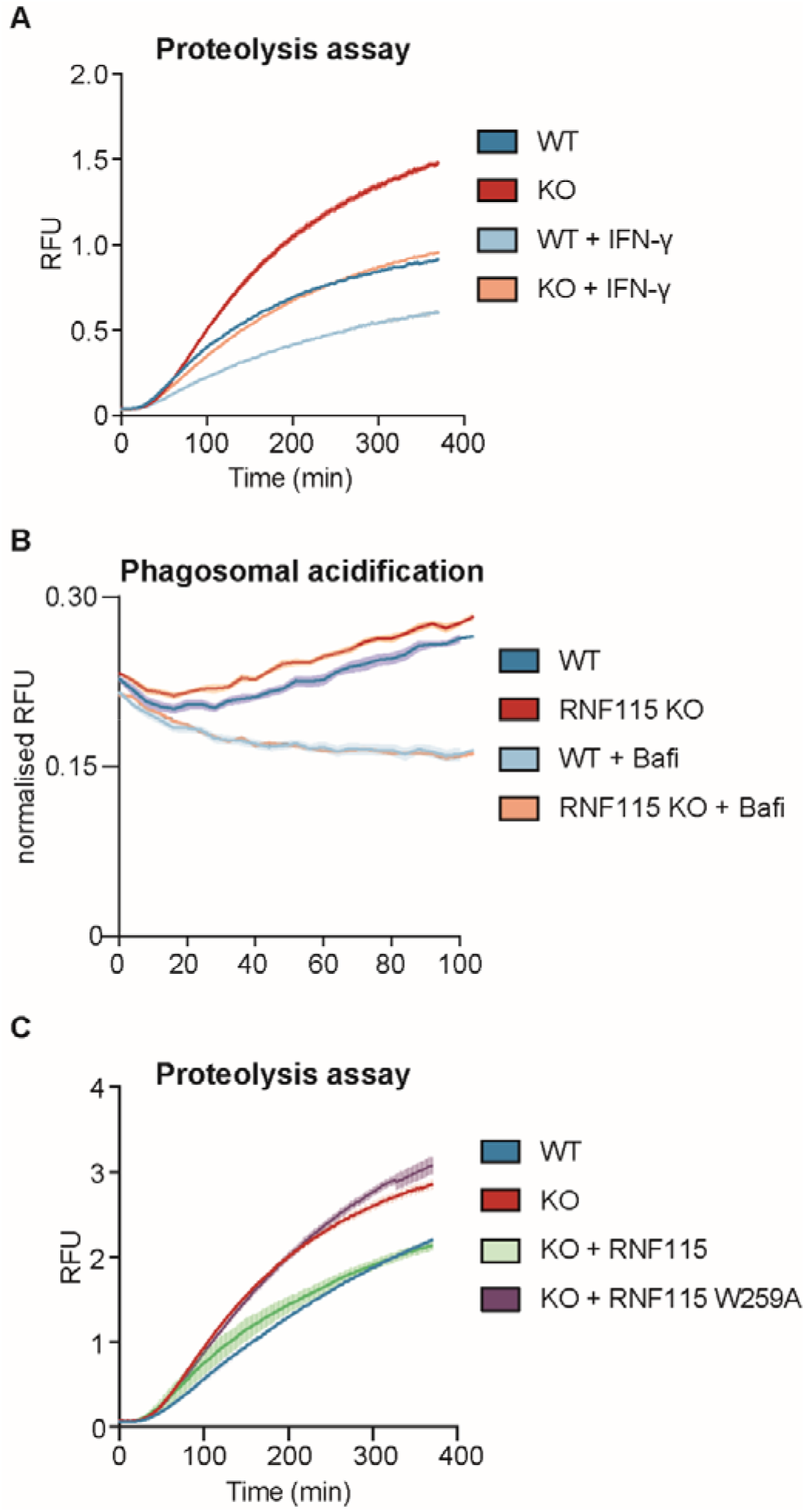
Loss of RNF115 increases phagosomal proteolysis and enhances innate immune responses to intraphagosomal bacteria. (A-B) Intraphagosomal proteolysis and acidification assays show that loss of RNF115 increases phagosome maturation in BMA cells. Bafi = Bafilomycin (C) Complementing the RNF115 KO with WT RNF115 or a W259A mutant which is unable to bind to E2 enzymes, shows that RNF115 ligase activity is required for the increase of phagosome maturation. Shaded areas represent SEM.

Altogether, these data indicate that RNF115 is a negative regulator of phagosomal maturation, and its ubiquitin ligase activity is required for its effect on phagosome maturation.

### Loss of RNF115 reduces ubiquitylation of phagosomal vesicle trafficking proteins and may affect vesicle fusion through VAMP8/Syntaxin7 SNARE complex

As our data indicated that the E3 ligase activity of RNF115 was required for the observed changes in phagosome maturation, we next performed a pull-down of ubiquitin remnant Gly-Gly peptides from phagosomal proteins to identify putative RNF115 substrates. Whilst these peptides can also originate from protein neddylation or ISGylation, previous studies have shown that 95% of all Gly-Gly peptides identified using this antibody enrichment approach arise from ubiquitylation (Kim et al., 2011). Moreover, total abundance of ubiquitin vs ISG15 in phagosomal extracts was about 60-fold higher, suggesting that ubiquitylation is in fact the primary modification, even following IFN-γ treatment. We performed Gly-Gly pulldowns from polystyrene bead phagosome extracts (^~^100 μg per channel) which was labelled with 16-plex tandem mass tags (TMT) from WT and RNF115 KO macrophage phagosomes in response to IFN-γ. We identified and quantified a total of 680 Gly-Gly (K) peptides from phagosomal proteins. Thirty-one Gly-Gly peptides showed increased abundance in RNF115 KO phagosomes (log2>0.6; p<0.05), whilst 131 peptides were significantly lower in the RNF115 KO compared to WT phagosomes (log2<-0.6; p<0.05) **(Figure 5A and Supplementary Table 4).**

**Figure 5:**
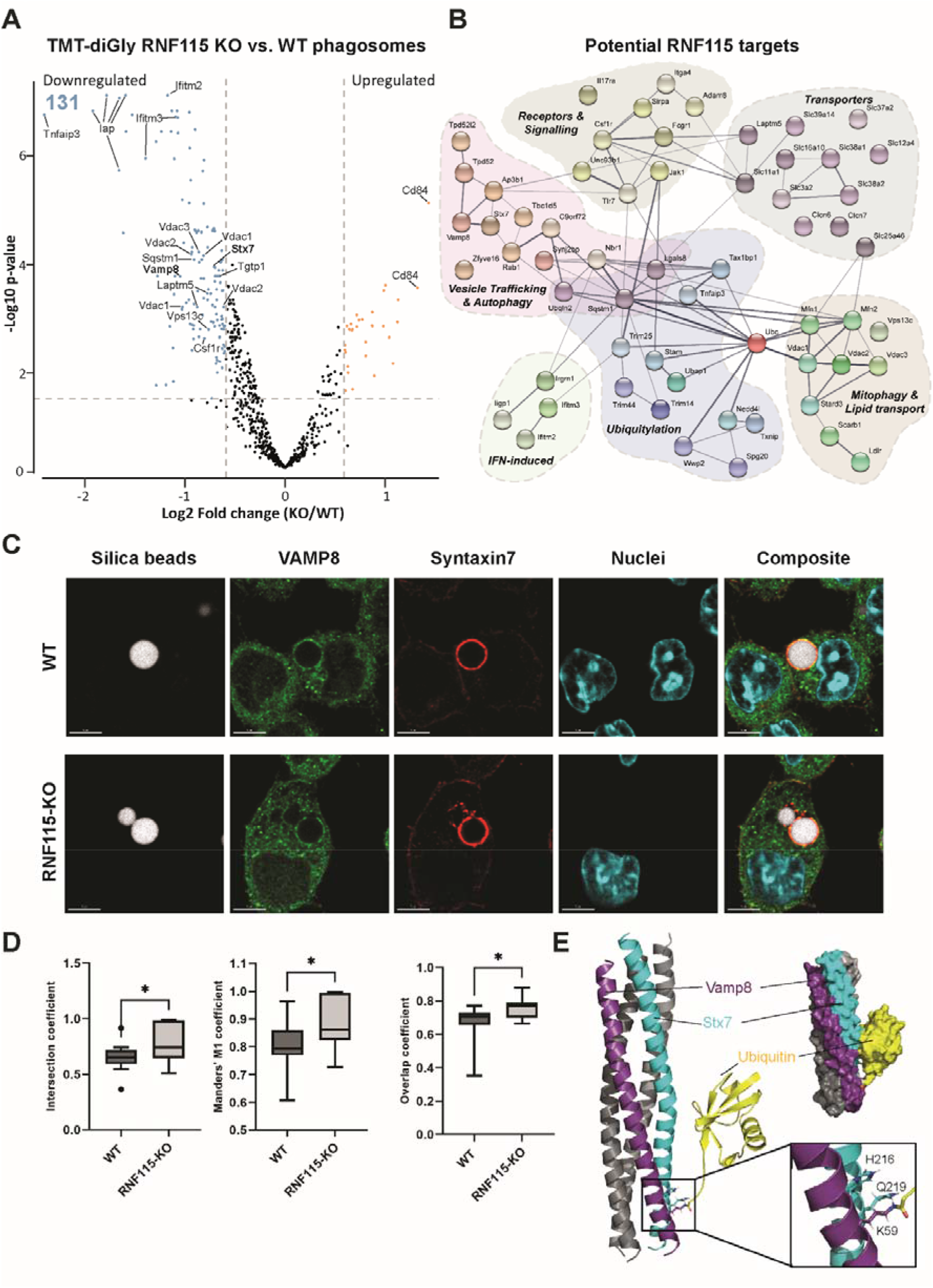
Pull-down of ubiquitylated proteins from phagosomes from WT and RNF115 KO macrophages indicates RNF115 ubiquitylates various proteins involved in immune responses and vesicle trafficking and may affect formation of the VAMP8/STX7 SNARE complex. (A) Volcano plot of proteomics experiment of Gly-Gly peptides from phagosomal extracts of WT and RNF115 KO macrophages. Selected proteins are highlighted. (B) Network representation (from STRING v11) (Szklarczyk et al., 2019) of selected proteins with reduced ubiquitylated peptide abundance in response to loss of RNF115, i.e. potential phagosomal substrates of RNF115. (C) Immunofluorescence micrographs of VAMP8 (green) and Syntaxin-7 (STX7; red) show strong co-localisation around 3 μm silica bead phagosomes in BMA macrophages. Nuclei are stained with Dapi. Scale bar is 5 μm. (D) Colocalization of VAMP8 (green) and STX7 (red) on individual phagosomes between WT and RNF115 KO cells represented by intersection, Manders’ M1 and overlap coefficients. E) Structure of VAMP8 and STX7 complex onto which a single ubiquitin molecule on VAMP8 K59 was modelled. Inlay: ubiquitylation of K59 disrupts a hydrogen bond between VAMP8 K59 and STX7 Q219.

Increased ubiquitylation sites upon loss of RNF115 included ubiquitin on K29, the NADPH-Oxidase complex member Cybb (K381/K567) and CD84 (SLAMF5) (K261) **(Supplementary Table 4).**

Proteins that showed lower levels of ubiquitylation in response to RNF115 KO were related to the innate immune response, signalling and receptors (IFITM2/3, IRGM1, JAK1, TLR7), mitophagy (Mfn1, VDAC1/2/3), amino acid, ion and sugar transporters, vesicle transport and autophagy (RAB1A, VAMP8, STX7) and ubiquitylation (STAM, SQSTM1, Ubiquitin K27 and K63, TNFAIP3/A20). Moreover, Tbc1d5, which may act as a GTPase-activating protein for Rab7a (Jimenez-Orgaz et al., 2018, Seaman et al., 2009, Seaman et al., 2018), was less ubiquitylated in RNF115 KO phagosomes, indicating that RNF115 may regulate Rab7a activity indirectly **(Figure 5B).** These proteins do not change in abundance between RNF115 KO and WT phagosomes **(Supplementary Table 4),** indicating RNF115-mediated ubiquitylation might not affect the degradation of these putative substrates, but rather affect protein or vesicular trafficking.

The ubiquitylation of the SNARE proteins VAMP8 and STX7 both decreased in RNF115 KO compared to WT phagosomes. Since these proteins are known to form a complex important for vesicle trafficking, we tested whether their ubiquitylation might affect their interaction and/or their function in vesicle fusion. Fluorescence microscopy validated high-co-localisation of VAMP8 and STX7 around the phagosome membrane **(Figure 5C).** Loss of RNF115 increased the co-localization slightly, but significantly, indicating that RNF115-induced ubiquitylation of this complex can regulate its trafficking **(Figure 5D).**

To test whether VAMP8/STX7 ubiquitylation affects SNARE complex formation, we performed structural modelling to predict if the two identified ubiquitylation sites (STX7 K138 and VAMP8 K59) would affect the structure and function of the SNARE complex **(Figure 5E).** While the STX7 K138 is in a disordered region, interrogation of the X-ray structure of the SNARE complex comprising STX7, VAMP8, STX8, and Vti1b [PBD identifier 1GL2] (Antonin et al., 2002) identified VAMP8 K59 as an integral part of the typical SNARE four-helix bundle. Interestingly, the amine side chain of VAMP8 K59 is connected via hydrogen bonding to residues H216 and Q219 of the adjacent STX7. A docked model of the VAMP8 K59 ubiquitylated SNARE complex suggests that ubiquitylation of the K59 side chain may disturb the local interface between STX7 and VAMP8. Nonetheless, based on the model, it appears that the ubiquitylation of VAMP8 K59 can be accommodated without unfolding of the SNARE four-helix bundle and is stabilised by salt bridges between STX7 and ubiquitin **(Figure 5E).** This indicates that the main structural and functional effect of VAMP8 K59 ubiquitylation is likely to be more indirect, for instance by affecting the formation and rearrangement of the SNARE complex, by recruitment of ubiquitin-binding proteins, or its membrane interactions.

### Loss of RNF115 promotes phagosome maturation and *Salmonella* adaptation and replication within macrophages

Given that loss of RNF115 promotes phagosome maturation, we next investigated whether RNF115 deficiency impacts survival of the Gram-negative bacterium *Salmonella* Typhimurium within macrophages. *Salmonella* Typhimurium is a facultative intracellular bacterium which is able to survive within the phagosome. Following invasion, Salmonella induces formation of a vacuole around the bacterium, called the Salmonella-containing vacuole (SCV), that allows for protection against host cytosolic antibacterial responses. For survival and replication inside phagocytes, *Salmonella* uses a second pathogenicity island (SPI-2) that is required for survival in the low pH of these cell types(Rappl et al., 2003, Srikanth et al., 2011). First, Phagocytosis assays revealed that RNF115 KO deficiency did not impact uptake of beads or any of the tested pathogens **(Supplementary Figure 4A-B).** We then used infection assays to assess whether *Salmonella* survival is affected by loss of RNF115. The analysis revealed that ablation of RNF115 facilitates *Salmonella* replication compared to WT, independent of IFN-γ **(Figure 6A).** Increased *Salmonella* survival corresponded with enhanced secretion of pro-inflammatory TNF-α and IL-6 *upon Salmonella* Typhimurium **(Figure 6B).** Moreover, we used two Salmonella reporters encoding stress-inducible promoters fused to an unstable GFP variant, which are able to report on quick changes in the *Salmonella* surrounding environment. Consistently, *Salmonella* displays higher SPI-2 activity within RNF115 KO cells suggesting faster intracellular adaptation and potentially replication **(Figure 6C).** We further identified that a greater percentage of Salmonella are exposed to acidic pH within macrophages lacking RNF115 compared to their WT counterpart **(Figure 6D).** These data suggest that loss of RNF115 leads to faster phagosome maturation which leads to enhanced *Salmonella* adaptation and replication within macrophages, but also induces increases innate immune responses to bacterial pathogens from the phagosome.

**Figure 6:**
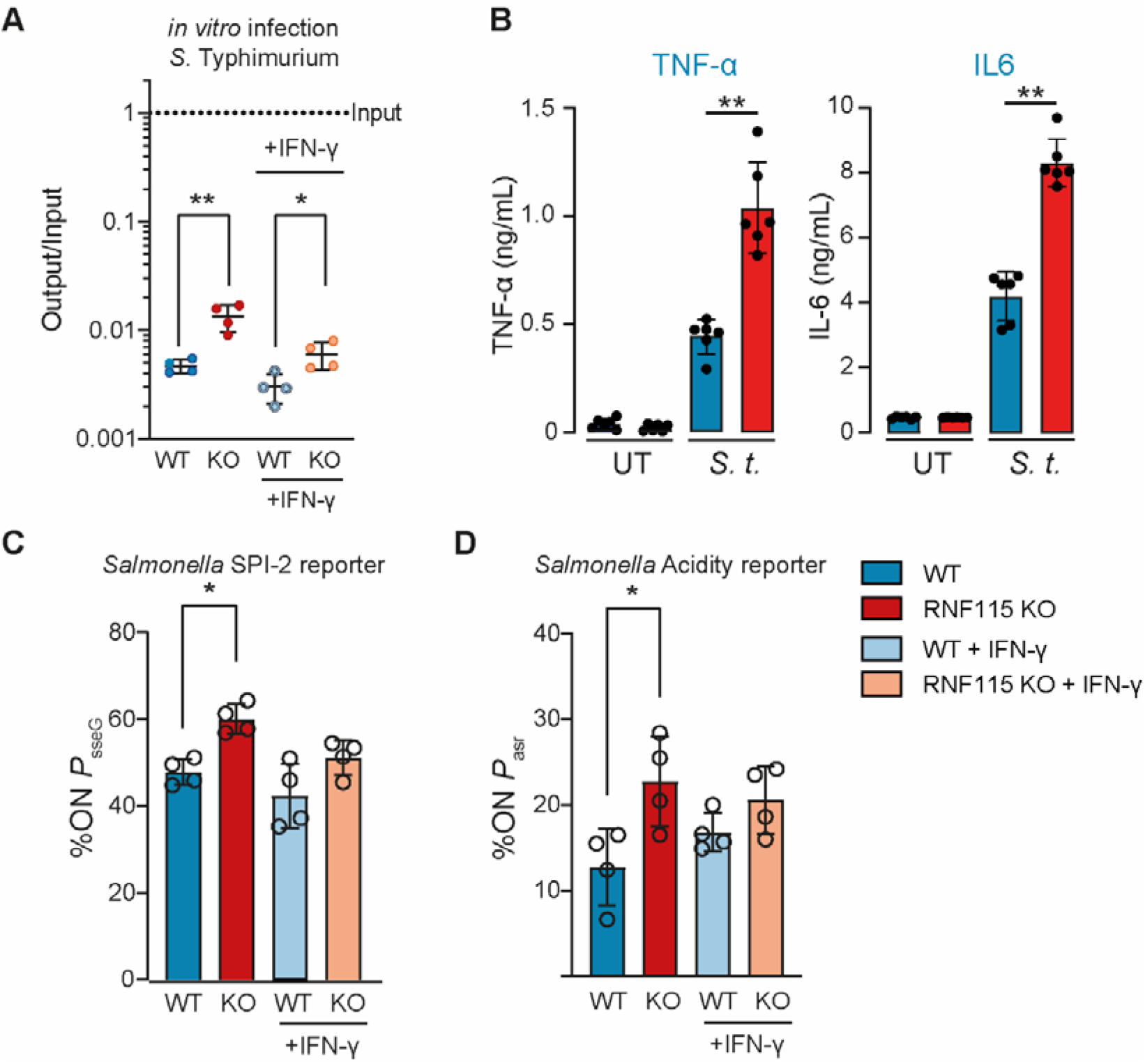
Loss of RNF115 promotes *Salmonella* adaptation and replication within macrophages and enhances innate immune responses. A) Infection of WT and RNF115 KO BMDMs with *S*. Typhimurium for 4 hrs shows an increased survival in RNF115 KO macrophages. (B) Infection with *Salmonella* Typhimurium for 6 hrs shows increased TNF-α and IL6 secretion in RNF115 KO BMDMs. Data of 5 independent replicates. MOI = 10. Error bars represent SEM. *=p<0.05; **=p<0.01; by paired two-tailed Student’s t-test. C-D) *Salmonella* SL1344 encoding stress-inducible promoter fusions from BMA cells infected for 4 hours display stronger SPI-2 response (*P*_sseG_) and acidity exposure (*P*_asr_) in RNF115 KO compared to WT cells. Each dot represents a biological replicate. Groups were compared using an unpaired two-tailed t-test. Error bars represent standard deviation. *, P < 0.05.

### Loss of RNF115 reduces tissue damage and inflammatory response to *S. aureus* infection *in vivo*

Next, we tested whether enhanced RNF115-mediated phagosome acidification impacts innate immune sensing of the Gram-positive *Staphylococcus aureus* which has been shown to be key for host defence (Ip et al., 2010). WT and RNF115 KO BMDMs were infected with *S. aureus* for 6 hrs and examined for cytokine release by ELISA. As a result, RNF115 KO macrophages indeed elicited higher levels of pro-inflammatory TNF-α and IL-6 cytokine secretion in response to *S. aureus* compared to WT **(Figure 7A).**

**Figure 7:**
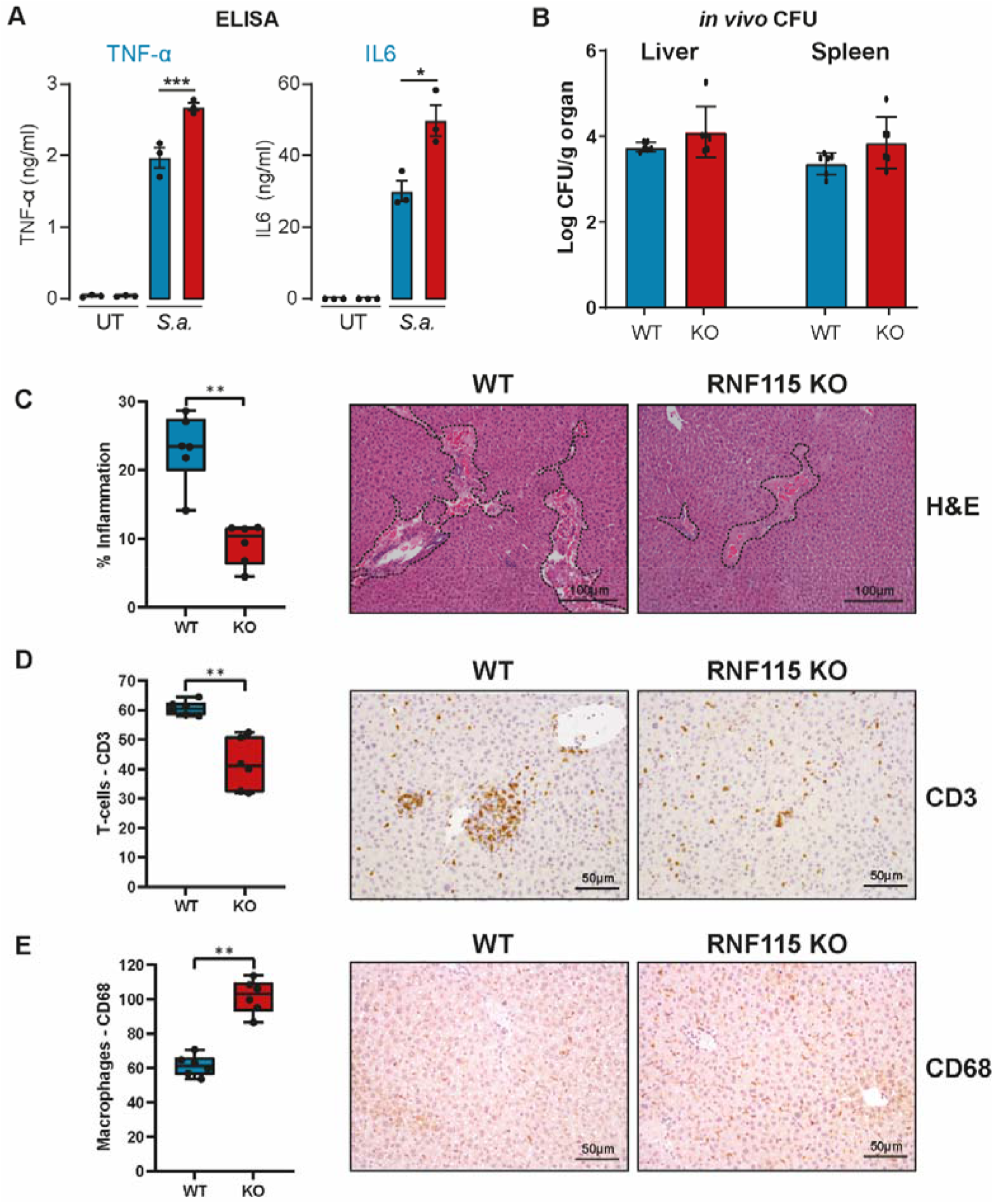
Loss of RNF115 increases inflammatory response *in vitro* but reduces tissue damage to *S. aureus* infection *in vivo*. (A) ELISA data show that loss of RNF115 increases TNF-α and IL6 secretion after 6 hrs of infection with *S. aureus*. Error bars represent SEM. *=p<0.05; ***=p<0.001; by paired two-tailed Student’s t-test. B) WT and RNF115 mice were infected with *S. aureus* and after 48 hrs CFUs were counted in liver and spleen. There was no significant difference between WT and RNF115 KO mice. C) Quantification of the hepatic lesion area in the livers from wild-type (WT) and RNF115 KO (KO) mice after 48 hours infection with *S. aureus*. Haematoxylin and eosin (H&E)-stained liver sections of one representative animal per genotype are shown on the right, dotted line depicts the damaged area. (D) Average cell counts per high power field (HPF) of CD3 positive T-cells, and E) CD68 positive macrophages in the livers from wild-type (WT) and RNF115 KO (KO) mice after 48 hours infection with *S. aureus*. Representative immunohistochemistry sections are shown on the right. Groups were compared using an unpaired two-tailed t-test. The statistical significance of the comparisons is indicated as follows: **, P < 0.01. Each dot represents an individual animal.

This data prompted us to test whether augmented bacterial sensing from phagosomes can lead to better host defence against *S. aureus in vivo*. For *in vivo* experiments, we infected six WT and RNF115 KO mice with *S. aureus*. RNF115 KO mice generated within this project have no obvious phenotype and breed with Mendelian ratios. After 48 hours, mice were humanely sacrificed, and livers and spleens were analysed for CFUs and by immunohistochemistry (IHC). Data showed that the loss of RNF115 did not significantly affect CFUs in livers and spleens of infected mice **(Figure 7B).** However, RNF115 KO mice showed significantly reduced infection/inflammation-dependent tissue damage **(Figure 7C)** and recruitment of CD3 T-cells **(Figure 7D).** However, RNF115 KO mice showed higher levels of CD68-stained macrophages **(Figure 7E)**(Wen et al., 2021), suggesting a higher level of tissue repair after infectious insult. These data indicate that loss of RNF115 reduces tissue damage in response to pathogenic bacteria *in vivo*.

## Discussion

Phagosomes are key organelles in innate immunity. As large intracellular vesicles, they are in continuous interaction with other vesicle trafficking pathways of the cell and in constant change. Here, we show for the first time that ubiquitylation plays a major role in regulating vesicle trafficking pathways to the phagosome and, thereby, regulating phagosome functions.

While ubiquitylation has been studied for many years in the endosomal system for its role in receptor sorting and multi-vesicular body (MVB) formation (Frankel & Audhya, 2018, Haglund & Dikic, 2012), and significant work has gone into characterising bacterial ubiquitin enzymes of intracellular pathogens such as *Legionella* (Vozandychova et al., 2021), little is known about the role of endogenous ubiquitylation on the phagosome. In 2005, it was shown that ubiquitin accumulates around the phagosome and that loss of E1 activity resulted in accumulation of Fc-receptors, suggesting a defect in the sorting of these receptors (Lee et al., 2005). Recently, it was shown that loss of the K63 specific E2 ligase Ubc13 (Ube2n) affects phagosome maturation leading to accumulation of apoptotic bodies in *C. elegans* (Liu et al., 2018). Furthermore, we have shown that receptor ubiquitylation can serve as a scaffold for pro-inflammatory signalling from the phagosome (Guo et al., 2019).

Our data indicates that phagosomes are not only rich in ubiquitylation, but they also contain significant amounts of atypical chains such as K27 and K33 chains whose biological function is poorly understood (Kulathu & Komander, 2012, van Huizen & Kikkert, 2019). It appears that K27 may also serve as a scaffold to recruit specific signalling pathways, as recent data showed that K27 ubiquitylation of BRAF by ITCH – which is also enriched on phagosomes – regulates MEK-ERK signalling (Yin et al., 2019). Interestingly, the UbiCRest experiment showed that most ubiquitylation on the phagosome is removed by using the K63-specific DUB AMSH-LP. This suggests that K63 chains may present the basis for chain architecture of polyubiquitylated phagosomal proteins. Future work will be required to characterise the roles of atypical chain types in phagosome biology.

It needs to be noted that IFN-γ activation also increases phagosomal ISGylation, the covalent addition of the interferon-induced ubiquitin-like modifier ISG15, which also leaves a Gly-Gly tag on proteins after tryptic digest. It is therefore possible that some of the sites identified were ISGylated rather than ubiquitylated. However, estimations of ubiquitin (8.6 kDa, log2 intensity 30) and ISG15 (17.8 kDa, log2 intensity of 25) abundance suggests that ubiquitin is about 60-fold more abundant on the phagosome than ISG15. This suggests that most of the Gly-Gly sites identified are in fact ubiquitylation sites. Chain-type specific TUBEs (Heap et al., 2017, Hjerpe et al., 2009) or the use of LB-Pro (Swatek et al., 2018, Swatek et al., 2019), an ISG15 specific protease, may provide more insights into the substrates of specific ubiquitin chain types in the future.

Nonetheless, our data provide a first glance at ubiquitylated proteins from the phagosome or in fact any organelle within the endo-lysosomal system. The diversity of substrates is astonishing, as is the fact that many ubiquitylation sites on functional domains of proteins, such as the SNARE domains of VAMP proteins, will likely disrupt protein-protein interactions. Further characterisation of the ubiquitylation of these proteins is hampered by the observation that antibodies often fail to bind recognition sequences once they are ubiquitylated. In this paper, for example, we show that ubiquitylated RAB7 is not detectable by Western Blot and that only after deubiquitylation, the protein appeared on the blot. The use of tandem ubiquitin binding entities (TUBEs) (Hjerpe et al., 2009) pulldowns with subsequent deubiquitylation may be a successful strategy to avoid difficulties.

We identified more than 30 E3 ligases on the phagosome. So far, only RNF19b (NKLAM) has been identified as an E3 ligase with the ability to regulate phagosome functions (Lawrence & Kornbluth, 2012, Lawrence & Kornbluth, 2018). Lawrence *et al* showed that loss of RNF19b led to reduced killing of phagocytosed *E. coli* as well as reduced inflammatory responses. Our data showed RNF19b to be unique to phagosomes of IFN-γ activated macrophages, supporting the findings of this work. In the future, it would be of interest to study the effects of loss of RNF19b on phagosome function and proteome.

In this paper, we followed up on the role of RNF115 (BCA2, Rabring7) on the phagosome. RNF115 was identified as a binding partner of Rab7 (Mizuno et al., 2003), involved in endosomal sorting of EGFR (Smith et al., 2013) and is highly expressed in invasive breast cancers (Burger et al., 2005). Recent work placed RNF115 as a regulator of inflammatory responses by showing that it may SUMOylate IκBα (Colomer-Lluch & Serra-Moreno, 2017) and that it may polyubiquitylate MAVS and MITA/STING (Zhang et al., 2020), two proteins important for detection of cytosolic viral RNA and DNA, respectively. Both MAVS and STING are present on phagosomes. While STING levels were only slightly reduced, MAVS was ^~^12-fold downregulated upon loss of RNF115. This contradicts the findings of Zhang *et al*, as one would expect increased MAVS and STING levels in the RNF115 KO. Moreover, total protein levels of STING and MAVS were not affected in total cell lysates of BMDMs (data not shown). Nonetheless, our findings suggest that RNF115 may somehow regulate MAVS, however, more likely through protein/vesicle trafficking from the mitochondria rather than direct K48 ubiquitylation.

Our data indicates, that RNF115 has a multitude of functions. The main differences that we detected in RNF115 KO macrophages was a dysregulation of innate immunity and vesicle trafficking pathways, suggesting that RNF115 ubiquitylates vesicle trafficking proteins. Similar to Zhang *et al*, who showed that loss of RNF115 affected immune responses to HSV-1 (Zhang et al., 2020), innate immune responses were affected in our experiments. However, in our model, this was only observed for signalling from pathogens within the phagosome. Whilst RNF115 may affect signalling pathways directly, it may also be possible that loss of RNF115 affects the time that innate immune receptors spend signalling from the phagosomal membrane by abolishing proper recycling and sorting of these immune receptors. More work will be needed to characterise the molecular targets of RNF115 to understand the exact function of this E3 ligase in innate immunity.

Despite having a clear phenotype *in vitro*, loss of RNF115 only mildly altered the survival of bacteria *in vivo* and *in vitro*. This is likely due to some redundancy amongst other E3 ligases and the hard-wiring of phagosomal functions such as maturation and acidification. Nonetheless, the data show that phagosomal ubiquitylation plays an important role in regulating phagosome biology. Pathogens such as *Salmonella* Typhimurium have evolved to withstand host defences and it may therefore not be surprising *that Salmonella* adapts well to increased acidification within the RNF115 KO phagosome. Future work on additional ubiquitylation enzymes will be required to further understand the role of ubiquitylation on the phagosome.

Overall, this work is the first in depth characterisation and quantification of ubiquitin on phagosomes. Our novel approach allowed an unbiased analysis of ubiquitin chains enriched on phagosomes and the identification of ubiquitylated phagosomal proteins. Our results demonstrate the importance of ubiquitylation in vesicle trafficking and indicate the regulatory function in immune signalling. We identified the E3 ubiquitin ligase, RNF115, as a new regulator of phagosomal maturation. We show that its E3 ligase activity is essential for its ability to affect phagosomal proteolysis and it is a negative regulator of pro-inflammatory cytokine induction from the phagosome upon bacterial infection.

## Materials and Methods

### Cell culture

The murine macrophage cell line RAW264.7 was obtained from ATCC (#TIB-71). The immortalized mouse macrophage cell line BMA3.1A7 (accession: CVCL_IW58) was kindly provided by Kenneth Rock (Dana Farber Center, Boston, US) (Kovacsovics-Bankowski & Rock, 1995). Both cell lines were maintained in Dulbecco’s modified eagle medium (DMEM), 10 % (v/v) heat-inactivated foetal bovine serum (FBS), 1 mM L-Glutamine, 100 U/ml penicillin, 100 μg/ml streptomycin. Cells were maintained in this media under 5% CO_2_ at 37°C in a water-saturated incubator. Cells were tested every 3 months for mycoplasma infection but are not authenticated.

### Mice

Wild-type C57BL/6J and C57BL/6NTac mice were obtained from Charles River and Taconic, respectively. RNF115 KO (RNF115-DEL562INS28-EM1-B6N) C57BL/6NTac mice were kindly provided by the Mary Lyon Centre, MRC Harwell, UK. The mice were phenotyped by the Mary Lyon Centre and the data is available here: https://www.mousephenotype.org/data/genes/MGI:1915095. Mice were bred under appropriate UK home office project licenses.

### Isolation and culturing of bone-marrow derived macrophages

Bone marrow cells were collected from femurs and tibia of both male and female 6-to 8-week old C57BL/6J or C57BL/6NTac wild-type mice. Collected cells were treated with red blood cell lysis buffer (155 mM NH_4_Cl, 12 mM NaHCO_3_, 0.1 mM EDTA) and plated on untreated 10 cm cell culture dishes (BD Biosciences) in IMDM (Gibco) containing 10% heat-inactivated FBS, 100 units/ml penicillin/streptomycin (Gibco) and 15% L929 conditioned supplement. After 24 h, the cells in supernatant were transferred to untreated 10 cm Petri dishes (BD Biosciences) for 7 days for the differentiation into bone marrow-derived macrophages (BMDMs) (Heap et al., 2021).

### Antibodies

The following antibodies were purchased from Cell Signaling technologies: Vimentin (#5741). EEA1 (#2411), Rab7 (#9367), iNOS (#2977). Antibodies purchased from Abcam were the following: Cathepsin D (ab75852), RNF115 (ab187642) and Histone H3 (ab176842). Total-ubiquitin antibody was purchased from DAKO (ZO458). Anti-Syntaxin 7 (sc-514017) and anti-Vamp8 Alex Fluor 488-conjugate were purchased from Santa Cruz Biotechnology (sc-166820 AF488). Ubiquitin chain-specific antibodies K63 (APU3.A8), K11 (2A3/2E6), K48 (Apu.2.07) Ubiquitin were a kind gift from Genentech. Tubulin (GTX628802) was purchased from Genetex, Flag-M2 (F3165) from Sigma Aldrich. RNF115 antibodies were produced in sheep by the antibody group of the Division of Signal Transduction Therapy (DSTT), MRC-PPU, University of Dundee and are available through https://mrcppureagents.dundee.ac.uk/.

### Proteins

All proteins in this work were kindly provided by Dr Axel Knebel, Protein Purification and Assay Development Team (PPAD), MRC-PPU or by the Protein Purification Team headed by James Hastie, DSTT, MRC-PPU. E3 ligases produced were RNF115 and RNF115 W259A. Deubiquitylase enzymes produced were: Otulin (Q96BN8), AMSH-LP (Q96FJ0), USP2 (NP_741994), vOTU (3ZNH_A), Cezanne (Q6GQQ9), TRABID (Q9UGI0). The Tandem Ubiquitin Binding Entity (TUBE) was HALO-Ubiquilin/NZF2 (Q9UMX0). Proteins were expressed as His6-tagged fusion proteins in BL21 (DE3) *E. coli* followed by a PreScission proteinase cleavage tag.

### Phagosome isolation

Phagosomes were isolated according to previous methods (Hartlova et al., 2017, Trost et al., 2009). In short, carboxylated polystyrene beads of 0.8 μm (Estapor/Merck) were internalised by macrophages for 30 minutes at 37°C, 5% CO_2_. For chase experiment cells were washed with pre-warmed PBS and fresh growth media for additional incubation times. After incubation, cells were washed in ice-cold PBS and scraped into 50 ml Falcon tubes while kept on ice. Non-internalized beads were removed, and cells were re-suspended in 1 ml of HB homogenization buffer (8.55% (w/w) sucrose, 2.5 mM imidazole, pH 7.4) supplemented with inhibitors of proteases (Complete, Roche) and phosphatases (1.15 mM sodium molybdate, 4 mM sodium tartrate dihydrate, 1 mM freshly prepared sodium orthovanadate, 5 mM glycerophosphate, all Sigma), and 100 mM N-Ethylmaleimide (NEM) to block DUBs. Cells were homogenized and mixed with 68% sucrose solution (68% (w/w) sucrose, 2.5 mM imidazole, pH 7.4). A sucrose gradient was prepared by layering 68%, lysate, 35%, 25% and 10% sucrose solution. The gradients were centrifuged in an ultracentrifuge (Beckman Coulter) at 24,000 rpm (72,300 ×g) for 1 hour at 4 °C. After the centrifugation, the latex beads containing phagosomes were visible by showing a blue band between the 10% and the 25% sucrose layers. The phagosomes were collected into new ultracentrifuge tubes using a thin-tipped transfer pipette. Phagosomes from two gradients were combined into one ultracentrifuge tube and were then washed by adding cold PBS until the volume reaches 1 cm below the top of the tube. After mixing, phagosomes were pelleted by centrifugation at 15,000 rpm (28,400 ×g) for 15 min at 4°C. After removing the supernatant, the phagosomes were either used for experiments or stored at −80°C.

### Phagosome functional assays

The fluorogenic assays for phagosomal proteolysis and acidification were adapted from Russell laboratory (Yates et al., 2009). Cells were seeded into a 96 well plate at 1x 10^5^ cells per ml 24 h prior to the experiment. Carboxylate silica beads (3 μm, Kisker Biotech) were conjugated with DQ red BSA (Molecular Probes) or pHrodo (Molecular Probes) and incubated for 3 minutes at 1:100 in binding buffer (1% FBS in PBS pH 7.5) with seeded macrophages at 37°C. Solution was replaced with warm binding buffer and cells were immediately measured at 37°C. Real-time fluorescence was measured using a SpectraMax Gemini EM Fluorescence Microplate Reader (Molecular Devices), set as maximal readings per well to allow reading time intervals of 2 min. The excitation/emission wavelengths were 590/620 (DQ red BSA), 650/665 (Alexa Fluor 640), 560/585 (pHrodo) for the proteolysis or acidification assay and measured in relative fluorescence units (RFU). Plots were generated from the ratios of signal/control fluorescence. Error bars were generated from standard error of the mean of 6 replicates.

### Phagocytosis assay

Cells were seeded into a 96 well plate at 1x 10^5^ cells per ml 24 h prior to the experiment. Alexa Fluor 488 BSA-coated silica beads (1 μm, Kisker Biotech) were incubated at 1:100 dilution for 60 minutes at 37°C. 100 μl trypan blue per well was used to quench signal of non-internalized particles. After removing trypan blue cells were measured using SpectraMax Gemini EM Fluorescence Microplate Reader (Molecular Devices), set at excitation/emission wavelengths 495/519 nm and measured in relative fluorescence units (RFU).

### Coupling HALO-Ubiquilin/NZF_2_ to HALO resin

200 μl packed HaloLink resin (Promega) was washed three times in HALO-binding buffer (50 mM Tris, 150 mM NaCl, 0.05% NP-40, 1 mM DTT). Beads were incubated with 820 μg HALO-Ubiquilin/NZF_2_ in 1 ml HALO-binding buffer for 16 h at 4°C in a rotating wheel. Beads were washed three times in HALO-binding buffer and used immediately or stored up to two weeks at 4°C in HALO-binding buffer supplemented with 0.01% sodium azide.

### HALO-Tab2 or Ubiquilin/NZF_2_ pull-down assay

To capture ubiquitin chains from cell extracts 10 μl of packed beads were incubated with 1mg cell extract protein. For enrichment of ubiquitylated proteins from phagosomal proteins after magnetic beads isolation, 60 μg phagosomal extract was incubated with 10 μl of packed beads. Samples were incubated for 16 h at 4°C at end-over-end mixing and beads were washed 3 times with IP washing buffer 50 mM Tris, 150 mM NaCl, 1% Triton-X 100. If no other treatment was required slurry was denatured by LDS sample buffer. Beads were transferred to SpinX tubes (Corning) to collect sample for SDS/PAGE.

### Immunoblot analysis

Phagosomes or cells were lysed directly in 2x Laemmli buffer (with 5% β-mercaptoethanol) and subjected to SDS-Page using 4-12% NuPAGE gels (Invitrogen) and immunoblotted to PVDF membrane. Membranes were blocked for 1 h at RT in 5% (w/v) skim milk in TBS-T (0.1% Tween-20) and subsequently incubated with primary antibodies overnight at 4°C. Horseradish-peroxidase (HRP) conjugated secondary antibodies were incubated with membranes, after which proteins were detected using ECL and X-ray films. Immunoblots were quantified using ImageJ software.

### Phagocytosis assay for immunofluorescence microscopy

Cells were seeded into wells containing No. 1.5H coverslips (Marienfeld-Superior) and left to adhere overnight. Cells were then treated with 20 ng/mL IFN-γ for 16 hours before the addition of Alexa Fluor-594-coated 3 μm silica beads diluted 1:2500 in normal medium for 30 minutes to allow for phagocytosis. Following this, cells were washed in PBS and fixed in ice cold methanol for 15 minutes at 4°C. Methanol was removed with sufficient washing in PBS, followed by permeabilization using 0.1% Triton X-100 for 10 minutes, and a blocking step in 5% BSA in PBS for 1 hour at room temperature. Anti-Syntaxin 7 (Santa Cruz Biotechnology: sc-514017) was diluted 1:10 in 5% BSA in PBS and incubated overnight at 4°C, followed by incubation with 1:1000 rabbit anti-mouse Alexa Fluor 633 (ThermoFisher Scientific). Anti-Vamp8 Alex Fluor 488-conjugate (Santa Cruz Biotechnology: sc-166820 AF488) was diluted 1:20 in 5% BSA in PBS and incubated overnight at 4°C. Cells were also stained with DAPI prior to mounting in ProLong Glass (ThermoFisher Scientific). Samples were imaged using a Zeiss LSM 800 in Airyscan mode with post-acquisition Airyscan processing performed in Zen Blue software. All imaging was performed at 63X magnification using identical zoom and voltage settings. Cropped image sections were generated using Imaris software.

### Co-localisation analysis

Co-localisation analysis was performed using Huygens software by first 3D-cropping images and defining regions of interest corresponding to Syntaxin7 and Vamp8 positive phagosomes. Automatic thresholding was performed using Costes method for statistical significance. Co-localisation coefficients between Syntaxin7 and Vamp8 were generated for individual phagosomes across 2 independent experiments.

### CRISPR/Cas9 BMA line production

Using the CRISPR/Cas9 genome editing system was performed with gRNAs in the BMA cell line targeting exon 1 of RNF115. Transfection was carried out using Fugene HD (Promega) reagent and 1 μg of gRNA and Cas9 D10A. Transfection was carried out overnight after which selection was induced using puromycin (1μg/ml) for 48 h. Cells were subcloned to maintain a monoclonal knockout population. Clones were validated using sequencing, immunoblotting and immunoprecipitation. BMA3.1A7 cell line was used for this method as it allowed CRISPR-Cas9 modification while this was not possible in RAW264.7 cells.

### E2 enzyme screening assay

WT RNF115 (0.4 μM) or RNF115 mutant (W259A) (0.4 μM) were incubated with 0.2 μM UBE1, 1 μM of indicated E2 enzyme and 5 μM of FLAG-ubiquitin in ubiquitin assay buffer containing 2 mM ATP in a total reaction volume of 30 μl. Samples were incubated at 30°C for 1 h while shaking 1000 rpm after which the reaction was terminated by the addition of LDS. The samples were subjected to SDS/PAGE and the formation of ubiquitin chains by RNF115 was assessed by immunoblotting with anti-FLAG.

### Deubiquitylase assay with endogenous proteins

Phagosomes were lysed in lysis buffer (50 mM Tris, 150 mM NaCl, 1% Triton-X 100, 5 mM DTT) and incubated with deubiquitylating enzymes at 30°C for 1 hour. Enzyme concentrations unless otherwise stated were: USP2 (2 μM), vOTU (2 μM), TRABID (5 μM), Cezanne (5 μM), AMSH-LP (2 μM), Otulin (5 μM). To stop reaction 1x LDS (Invitrogen) was added and immunoblotting was conducted.

### Cell lysis and immunoprecipitation

Cells were washed in ice-cold PBS two times after which they were lysed on ice in lysis buffer (50 mM Tris-HCl pH 7.5, 150 mM NaCl, 1% Triton-X 100, 1x Complete^™^ protease inhibitor cocktail tablet (Roche), 0.1 mM EDTA, 0,1 mM EGTA, 50 mM NaF, 5 mM sodium pyrophosphate, 1 mM sodium orthovanadate, 10 mM sodium b-glycerophosphate, 100 mM N-ethylmaleimide (NEM). After lysis in appropriate volume samples were incubated on ice for 20 minutes and then centrifuged at 350 xg for 20 min 4°C. Supernatant was acquired and protein concentration was determined according to the Bradford method using Protein Assay Dye Concentrate.

Required volume of protein G-Sepharose beads were washed three times with PBS before addition of antibody at 1:1 ratio. The final volume was adjusted with PBS to give 2:1 ratio to ensure sufficient mixing of beads. Beads were shaken for 2 hours at 4°C and then washed 2 more times with PBS. For 15 μl of protein G-Sepharose beads 1 mg protein lysate was used and incubated for 4h while shaking at 4°C. Samples were washed 3 times in washing buffer (50 mM Tris-HCl pH 7.5, 150 mM NaCl, 1% Triton-X 100). Finally, beads were resuspended in 1x Laemmli buffer and used for immunoblotting.

### Immunofluorescence

BMDM cells were seeded on coverslips 24 h before experiment and activated with IFN-γ, if needed. Carboxylate silica beads (3 μm, Kisker Biotech) were added at 1:1000 dilution for 30 minutes after which slides were washed with PBS 3 times. 4% PFA was added at RT 20 minutes, after which cells were permeabilized by 0.1 % Triton-X 100 in PBS for 5 min RT. Blocking was done in 5% BSA in PBS for 30 min at RT after which primary antibodies were added RNF115 (1:200) o/n at 4°C. After washing 3 times Alexa Fluor 594 coupled anti-rabbit (Life Technologies) was added at a dilution of 1:500 for 1h at RT. DAPI was used to counterstain DNA and slides were mounted in Mowiol.

### qPCR

BMA cells were seeded at a final concentration of 0.2 x 10^6^ into a 12-well plate. After stimulation, total RNA was extracted using RNeasy Purification Kit (Qiagen, Cat. No. 74104). Using iScript cDNA synthesis kit form Bio-Rad (170-8891) 0.5 μg RNA was reverse transcribed. PCR was performed in a 384 well plate and each 10 μl reaction contained 0.5 % cDNA, 0.5 μM primers and 5 μl SSo Fast EvaGreen Supermix from Bio-Rad (172-5204) according to manufacturer’s instruction on a CFX384 machine (Bio-Rad). Normalisation of mRNA was done to 18S and the relative gene expression levels, in comparison of control (unstimulated cells (UT)), were calculated according to the comparative cycle threshold. Levels of mouse RS18, IL −6 and TNFα mRNA were detected by validated QuantiTect primer assays 144 (Qiagen).

### Bacterial strains and culture conditions

All experiments with *Listeria monocytogenes* were performed with strain EDGe (original stock from Hao Shen, University of Pennsylvania) and its *hly* (EJL1) derivative were grown in brain heart infusion (BHI) broth. *Staphylococcus aureus* RN6390 (kind gift from Tracy Palmer, Newcastle University) were grown in Tryptone Soy Broth (TSB). Bacteria were incubated at 37°C with constant rotation. *Salmonella enterica sv* Typhimurium SL1344 and two *Salmonella* reporter strains encoding stress-inducible promoters fused to an unstable GFP variant for acidity exposure (*P*_asr_) and SPI-2 response (*P*_sseG_) were kindly provided by Dirk Bumann (Biozentrum Basel). *Salmonella* strains were grown in Luria-Bertani (LB) broth. All bacteria were used at mid-exponential phase for infection experiments, washed in ice cold PBS twice and, subsequently, re-suspended in cell culture media at the desired MOI (10 for *S*. Typhimurium, 25 for *Listeria* and 25 for *S. aureus*).

### Bacterial infection

Bacterial cultures were grown to mid-exponential phase and pelleted by spinning at 7,000 ×g for 5 min, washed in PBS and subsequently resuspended in DMEM containing 10% FBS. Bacteria was added at MOI 10 to BMA cells for 30 minutes. Afterwards cells were washed and resuspended in DMEM containing 10% FBS and 1.5 μg/ml (*Salmonella*) or 5 mM (*S. aureus* and *Listeria*) gentamycin for 30 min. Once again media was removed cells washed and resuspended in DMEM containing 10% FBS for indicated time points after which cells were lysed as needed.

CFUs were determined by lysing cells at indicated time points in 0.1% Triton-X 100 in PBS from which serial dilutions were produced and plated for o/n incubation at 37°C. On the next day bacterial colonies were counted and CFU was determined.

### Flow Cytometry for *Salmonella* reporters

The *Salmonella* biosensors strains used in this study carried low-copy episomal pSC101-derivatives with inducible gfp-ova fused to candidate promoters (Pasr–acidity, PssaG–SPI2). *Salmonella* from infected macrophages was fixed for 15 min in 4% PFA/PBS before analysis on a flow cytometer (Symphony, BD Biosciences) using thresholds on side scatter (SSC) to exclude electronic noise. The percentage of GFP positive Salmonella was determined using FlowJo software.

### *In vivo* mouse infection

Approval to conduct research on animals was granted by the United Kingdom Home Office (PC123A338) and Newcastle University. Adult wild-type and RNF115 KO C57BL/6NTac mice were fed a standard rodent diet and kept under controlled environmental conditions (12-h light-dark cycle). Mice were randomized. Both groups contained similar numbers of male and female mice. 6 mice of each genotype were infected, 6 mice of each genotype were injected with PBS as control. Previous experiments indicated that this number was reaching enough statistical power.

*S. aureus* RN6390 was cultured in TSB until an optical density of 0.8-1.0 at 600 nm was reached. The cells were harvested by centrifugation (10,000 ×g, 2 min) and washed twice with PBS pH 7.4. *S. aureus* RN6390 cells were resuspended in PBS pH 7.4 and used for infections. Each animal received 2 x 10^6^ CFU of *S. aureus* RN6390 in 0.2 ml of PBS pH 7.4 via intraperitoneal infection. Cell counts of the inoculum were verified by serial dilution and plating on TSB agar plates. The plates were subsequently incubated at 37°C for 24 hours, after which CFU were determined.

The mice were euthanized 48 hours after the infection via cervical dislocation, and livers were harvested for immunohistochemistry. For immunohistochemistry, liver samples were fixed in 4% formaldehyde in PBS pH 6.5. Tissue samples were processed using automated procedures to impregnate and subsequently embed samples in paraffin wax.

### Immunohistochemistry

Formalin-fixed and paraffin-embedded liver sections were cut into 3-μm thick sections, deparaffinised in xylene and rehydrated in ethanol and water. The sections were stained with Mayer’s haematoxylin and eosin using standard protocols and inflammatory foci in the tissue section was counted at X10 magnification.

For immunohistochemical detection, endogenous peroxidase activity was blocked using hydrogen peroxide. Tissue sections were subjected to heat-induced antigen retrieval in EDTA, pH 8.0 for CD3 and sodium citrate buffer, pH 6.0 for CD68. Non-specific protein binding was blocked using 20% swine serum at room temperature for 30 minutes, and then antibodies specific to CD3 (dilution 1:100; MCA1477, Bio-Rad), and CD68 (dilution 1:200; OABB00472, Aviva Systems Biology) were incubated overnight at 4°C (Leslie et al., 2020). The following day slides were washed with PBS and then incubated with the appropriate biotinylated secondary antibody; swine anti-rabbit 1:200 (eo353 Dako) for CD68 or goat anti-rat 1:200 (STAR80B Serotec) for CD3. After washing, slides were then incubated with Vectastain Elite ABC Reagent (Vector Laboratories). Antigens were visualised using DAB peroxidase substrate kit (Dako) and counterstained with Mayer’s haematoxylin. The negative control was obtained by the replacement of primary antibody with PBS. A Nikon ECLIPSE Ni-U microscope (NIS-Elements Br, Nikon, UK) microscope was used for photographic material acquisition at 100x and 200x magnification. Irrespective of intensity, the percentages of positive cells *versus* total cell number were calculated using fifteen random fields per liver section in 6 mice per genotype. ImageJ (Fiji) was used for analysis as published previously (Marin-Rubio et al., 2019). Immunohistochemistry was analysed blinded by an expert, using an unpaired two-tailed t-test using GraphPad Prism (version 8.0.1).

### ELISA

Cells were seeded in a 96-well plate and infected with *S. aureus* or *L. monocytogenes* at a MOI of 10 or 25 (as indicated). The supernatant was collected at 6 h post-infection and centrifuged down. ELISA for TNF-α (DuoSet mouse ELISA kits from R&D Systems) were performed according to manufacturer’s instruction.

### Gly-Gly enrichment of ubiquitylated phagosomal peptides

PTMScan ubiquitin remnant motif (K-⍰-GG) kit (Cell Signalling Technology, cat. no. 5562) was used according to manufacturer’s instruction. All steps were performed in LoBind tubes. Phagosomal samples were prepared and digested as described above but desalting of samples was tC18 SepPak cartridge (Waters) as described by the manufacturer. K-⍰-GG beads enrichment of peptides was performed as described by (Udeshi et al., 2013). Briefly, K-⍰-GG beads were cross-linked in cross-linking buffer of 100 mM sodium borate (pH 9.0) with freshly added 20 mM DMP. Antibody was incubated with cross-linking buffer for 30 minutes at RT with gentle end-over-end rotation. Reaction was stopped by washing beads in antibody blocking buffer (200 mM ethanolamine). After washing beads were incubated in antibody blocking buffer for 2 hours at 4°C with gentle rotation. Cross-linked antibody was washed three times in ice-cold IAP buffer (50 mM MOPS (pH 7.2), 10 mM sodium phosphate and 50 mM NaCl). Dried down peptides were resuspended in ice-cold IAP buffer and centrifuged down to remove insoluble material. Supernatant was added to K-⍰-GG beads containing tubes (here to 300 μg phagosomal proteins add 5 μl packed beads). Samples were incubated for 1 h at 4°C with gentle end-over-end rotation. Samples were centrifuged down at 800 ×g for 1 min at 4°C and supernatant was removed. Beads were washed twice in IAP buffer followed by three washes in ice-cold PBS. For K-⍰-GG peptide elution after the final wash 50 μl of 0.15% (v/v) TFA was used gently mixed and incubated at RT for 5 minutes. Samples were centrifuged and supernatant was collected in fresh tubes. Elution step was repeated and collected supernatant was cleaned up with StageTip desalting columns (Thermo-Fisher) as described by the manufacturer and dried down in a Speed-Vac concentrator (Thermo-Scientific) resuspended in 20 μl L/C water containing 3% acetonitrile (MeCN) (Merck) and 0.1% FA. From this a 1:5 dilution was prepared and 4 μl was injected immediately into the mass spectrometer.

### Preparation of ubiquitin AQUA peptides

Concentrated stock of isotopically labelled internal standard (heavy) peptides (M1, M1ox, K6, K6ox, K11, K27, K29, K33, K48, K63) were purchased from Cell Signaling Technologies and Cambridge Research Biochemicals (Cleveland, United Kingdom). All peptides were stored at −80°C and working stock concentration was prepared of individual peptides at 25 pmol/μl in 2% (v/v) MeCN, 0.1 % (v/v) FA. Experimental mixture of all heavy peptides was prepared at 250 fmol/μl in 2% (v/v) MeCN, 0.1 % (v/v) FA.

### Absolute ubiquitin quantification by Parallel Reaction Monitoring

PRM analysis was performed on an Orbitrap Fusion or QExactive HF mass spectrometer (Thermo-Fisher Scientific) with an Easy-Spray source coupled to an Ultimate 3000 Rapid Separation LC System (Thermo-Fisher Scientific) as described before (Heunis et al., 2020). With a 5 μl full loop injection samples were loaded directly onto an EASY-Spray column (15 cm x 75 μm ID, PepMap C18, 3 μm particles, 100 Å pore size, Thermo-Fischer Scientific).

Separation of peptides was conducted by reverse phase chromatography at a flow rate of 1 μl/min (Solvent A 98% (v/v) H_2_O, 2% (v/v) MeCN, (v/v) 0.1% FA and solvent B was 98% (v/v) MeCN, 2% (v/v) H_2_O, 0.1% (v/v) FA). After LC injection peptides were resolved with an isocratic gradient of 0.1% of solvent B (10 minutes), then an increase of 0.1% to 25.5% of solvent B for over 41 minutes and finished with a 5 minute wash of 90% solvent B to wash off any contaminations. Orbitrap Fusion mass spectrometer was operated in “tMS2” targeted mode to detect ubiquitin peptides. Included m/z values were selected by quadrupole, with 4 m/z isolation window, with maximum injection time of 100 ms and a maximum AGC target of 5e4. HCD fragmentation was performed at 30% collision energy. MS/MS fragments were detected in the Orbitrap mass analyser at a 200 m/z resolution. Quantification of ubiquitylated peptides was done with Skyline (version 3.5.0.9191). To avoid interferences extracted ion chromatogram of MS/MS for precursor ion was adjusted manually.

### Sample preparation and mass spectrometry analysis

Phagosomal proteins from RAW264.7 cells were extracted in 1% sodium 3-[(2-methyl-2-undecyl-1,3-dioxolan-4-yl)methoxy]-1-propanesulfonate (commercially available as RapiGest, Waters) in 50 mM Tris (Sigma) pH 8.0, supplemented with 5 mM of tris(2-carboxyethyl)phosphine (TCEP). Samples were heated at 65°C for 5 minutes and afterwards alkylated with 10 mM iodoacetamide (Sigma) for 20 minutes in the dark. Dithiothreitol (DTT) of 20 mM was added for 20 minutes to quench alkylation. Protein concentrations were determined using EZQ protein quantitation kit (Molecular Probes). Samples were then diluted to 0.1% RapiGest in 50 mM Tris/HCL and digested using Trypsin Gold (Promega).

RapiGest was removed by adding trifluoroacetic acid (TFA, Sigma) to a final concentration of 1%, shaking the samples at 37°C for 1 h and spinning them at 14,000 xg for 30 min. Peptides were desalted by solid phase extraction (SPE) using C-18 Micro Spin columns as described by the manufacture’s instruction (The Nest Group). Samples eluted from Micro Spin columns were dried down in SpeedVac concentrator (Thermo Scientific) and if needed stored at −80°C.

Phagosomes from BMDMs were lysed in 5% SDS in 50 mM TEAB pH 7.5. Ten μg protein was processed for proteomic analysis using the suspension trapping (S-Trap) sample preparation method (ProtiFi) as previously described (Heap et al., 2021). Samples were dried and stored at −80°C.

### Mass spectrometry analysis

For mass spectrometry analysis peptides were resuspended in HPLC-grade water containing 2% MeCN and 1% TFA to make a final concentration of 0.5 μg/μl. 4 μl of samples were injected for analysis. Peptides were separated using 50 cm Acclaim PepMap 100 analytical column (75 μm ID, 3 μm C18) in conjunction with a Pepmap trapping column (100μm × 2 cm, 5 μm C18) (Thermo Scientific) analysed with Orbitrap Fusion Tribrid mass spectrometer (Thermo-Fisher Scientific). A three-hour gradient was performed with 3 % solvent B to 35 % solvent B (solvent A: 3% MeCN, 0.1% FA; solvent B: 80% MeCN, 0.08% FA). Settings for data acquisition were MS1 with 120,000 resolution, scan range 400-1600, charge state 2-5, AGC target of 200,000 and dynamic exclusion of 60 ms with repeat count 1. Peptide ions were fragmented using HCD (35 % collision energy) with a resolution of 15,000, and AGC target of 50,000 with a maximum injection of 60 ms. The whole duty cycle was set to 2.5 s during which the instrument performed “top speed” analysis.

Peptides from BMDM phagosomes were dissolved in 2% MeCN containing 0.1% TFA, and each sample was independently analysed on an Orbitrap Fusion Lumos Tribrid mass spectrometer (Thermo Fisher Scientific), connected to an UltiMate 3000 RSLCnano System (Thermo Fisher Scientific). Peptides (1 μg) were injected on a PepMap 100 C18 LC trap column (300⍰μm ID⍰×⍰5 mm, 5⍰μm, 100⍰Å) followed by separation on an EASY-Spray nanoLC C18 column (75⍰μm ID ×⍰50 cm, 2⍰μm, 100⍰Å) at a flow rate of 300⍰nl/min. Solvent A was 0.1% FA and solvent B was 80% MeCN containing 0.1% FA. The gradient used for analysis of samples was as follows: solvent B was maintained at 3% for 5⍰min, followed by an increase from 3 to 35% B in 180⍰min, 35-90% B in 0.5⍰min, maintained at 90% B for 4⍰min, followed by a decrease to 3% B in 0.5⍰min and equilibration at 3% B for 10⍰min. The Orbitrap Fusion Lumos was operated in positive-ion data-dependent mode. The precursor ion scan was performed in the Orbitrap in the range of 400-1,600⍰m/z with a resolution of 120,000 at 200⍰m/z, an AGC target of 400,000 and an ion injection time of 50⍰ms. MS/MS spectra were acquired in the linear ion trap using Rapid scan mode after HCD fragmentation. An HCD collision energy of 30% was used, the AGC target was set to 10,000 and dynamic injection time mode was allowed. The number of MS/MS events between full scans was determined on-the-fly to maintain a 3⍰s fixed duty cycle.

### Proteome quantification

Label-free quantification was performed using MaxQuant (v.1.5.7.4 (IFN-γ experiment) or v.1.6.3.4 (RNF115 KO)) with the following modifications: fixed modification: carbamidomethyl (C); variable modifications oxidation (M), acetylation (protein N-terminus), Deamidation (NQ), Glu->pyro-Glu, Ubiquitylation (GG, LRGG); label-free quantitation with minimum ratio count 2; maximum 5 modifications per peptide, and 2 missed cleavages. Searches were conducted using a murine Uniprot-Trembl database (downloaded March 26 2016, 28,245 entries (IFN-γ experiment) or downloaded May 5 2019, 25,231 entries (RNF115 KO)) and a list of common contaminants. Identifications were filtered at a 1% false-discovery rate (FDR). Quantification used only razor and unique peptides with a minimum ratio count of 2. “Re-quantify” was enabled. “Match between runs” was used with alignment time window 20 min and match time window 0.7 min. LFQ intensities were used for data analyses.

### Data analysis

Statistical analyses of most data was performed in GraphPad prism v9.0.2. Statistical data analysis of proteomics data was performed in Perseus (v.1.5.1.1 or 1.6.6.0) (Tyanova et al., 2016) or in the R statistical programming language (https://www.r-project.org/) using the LIMMA (Ritchie et al., 2015) and DEqMS (Zhu et al., 2020) packages. Contaminants were removed from data set. Using LFQ Intensities from MaxQuant analysis protein ratios were generated, logarithmized and significance of changes was analysed by using Student t-test analysis (p<0.05). Imputation was used at standard settings. GO-term enrichment analysis was performed using DAVID GO (v. 6.7) (Jiao et al., 2012). Here significantly changed proteins (fold change >2, p<0.05) were analysed against the background of all identified proteins. Networks were retrieved from String Database (Szklarczyk et al., 2015).

### Data availability

The mass spectrometry proteomics data have been deposited to the ProteomeXchange Consortium (Deutsch et al., 2017) via the PRIDE partner repository (Perez-Riverol et al., 2019) with the data set identifier: PXD026843.

## Acknowledgements

MT dedicates this paper to Rolf Wehler who taught him scientific thinking. We would like to thank the DNA cloning, protein production, antibody production, DNA sequencing facility, tissue culture and mass spectrometry teams of the MRC Protein Phosphorylation and Ubiquitylation Unit for their support. We would like to thank the Newcastle University Comparative Biology Centre and the Bioimaging Facility for support. We would like to thank Helen Wang and Tracy Palmer for providing bacterial strains, Axel Knebel and James Hastie for protein production and Thomas Macartney for support with the CRISPR work. We would like to thank MRC Mary Lyon Centre for providing the RNF115 KO mouse. We would like to thank Helen Walden for identifying the W259A mutation of RNF115. We would like to thank Dirk Bumann (Basel Biozentrum) for allowing us to use the *Salmonella* reporter strains. This work was funded by Medical Research Council UK (MC_UU_12016/5), Newcastle University start-up funding and a Wellcome Trust Investigator Award to MT (215542/Z/19/Z). AH is funded by the Knut and Alice Wallenberg Foundation. FRC is a Marie Sklodowska Curie Fellow within the European Union’s Horizon 2020 research and innovation programme under the Marie Skłodowska-Curie grant agreement No. 892252. FO is funded by Medical Research Council program grants MR/K0019494/1 and MR/R023026/1.

## Author contributions

OBG, TH and AH performed most experiments; JP, JLMR, DF, FRC, JI, BR and FO performed additional experiments; RS performed structural modelling; OBG, TH, JP and MT performed data analyses; MT and AH designed experiments and acquired funding; MT, OBG and AH wrote the paper with contributions of all authors.

