## Supplementary Figures for "The E3 ubiquitin ligase RNF115 regulates phagosome maturation and host response to bacterial infection"

Supplementary Figure 1

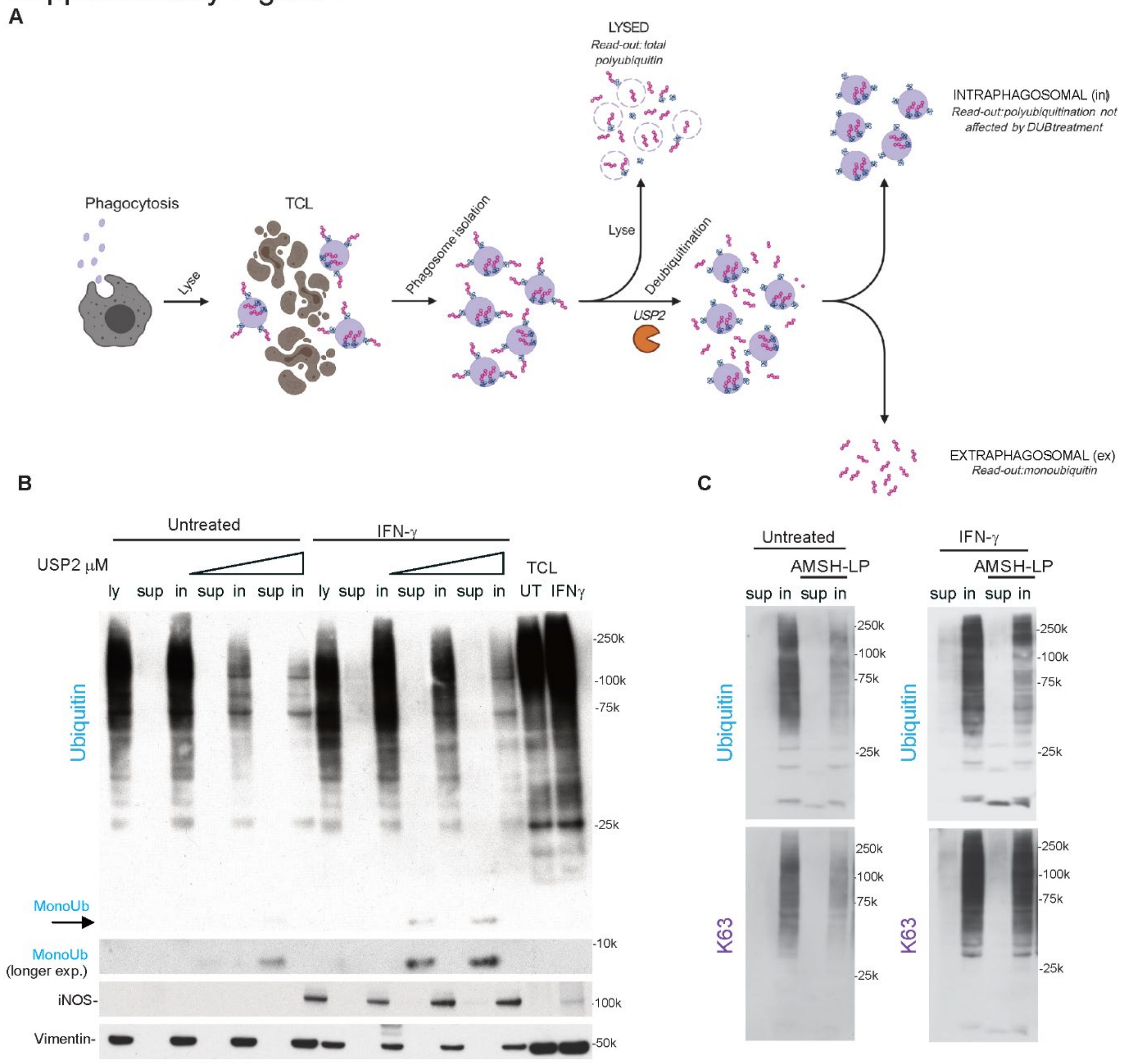


**Supplementary Figure 1: Phagosomal ubiquitylation is almost entirely cytoplasmic.** (A) Workflow: phagosomes were isolated and then either lysed using a detergent containing buffer or treated with increasing concentrations of (chain unspecific) USP2 to cleave off cytoplasmic ubiquitin chains. Phagosomes were then pelleted and the supernatant or the lysed pellet was analysed by Western Blot against ubiquitin. (B) Western blot of phagosomes treated as described in A). The data show that increasing amounts of USP2 can remove ubiquitin chains from the cytoplasmic side, leaving mono ubiquitin in the supernatant (sup). Intact phagosomes (in) show reducing amounts of polyubiquitin at the same time. Lysates (ly) serve as a control. iNOS serves as a control for IFN-activation. Vimentin serves as an attempt to control loading. (C) The same treatment of phagosomes with the K63-specific AMSH-LP shows that K63 chains are highly abundant on the cytoplasmic side of phagosomes. Western blots representative of two replicates.

### Supplementary Figure 2

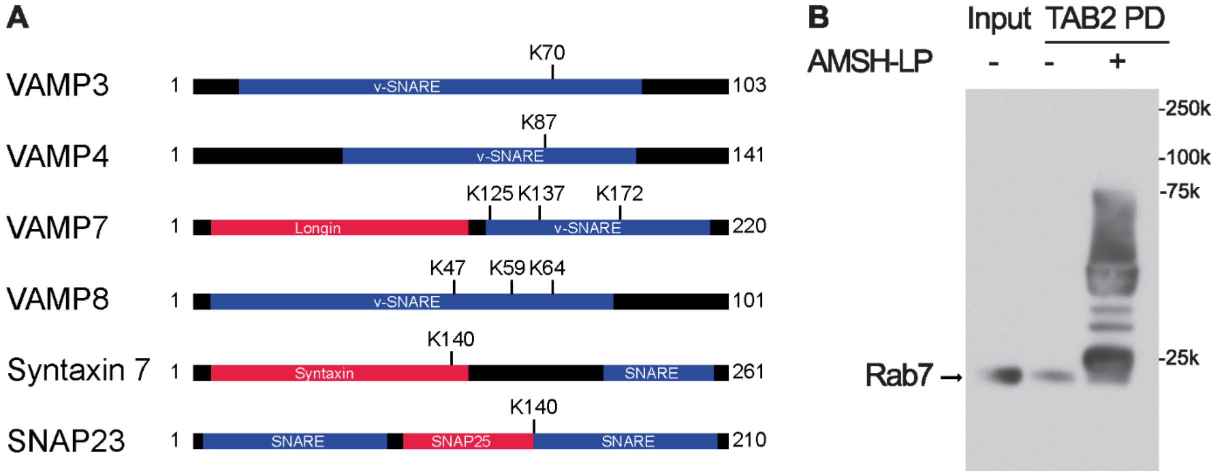


**Supplementary Figure 2: Ubiquitylation of vesicle trafficking proteins and TAB2-TUBE pulldown of ubiquitylated phagosomal proteins.** (A) Ubiquitylation sites of phagosomal SNARE proteins are almost entirely within SNARE domains, thereby blocking SNARE protein interactions. (B) Ubiquitylation of proteins affects their detection by antibodies. As an example, Rab7 is shown. TAB2-NFZ TUBE pulldown (PD) of K63 polyubiquitylated proteins and subsequent Western blot of Rab7 shows no ubiquitylated forms of Rab7 (probably by blocking of antibody antigen). Upon AMSH-LP treatment multiple forms of ubiquitylated Rab7 appear.

#### Supplementary Figure 3

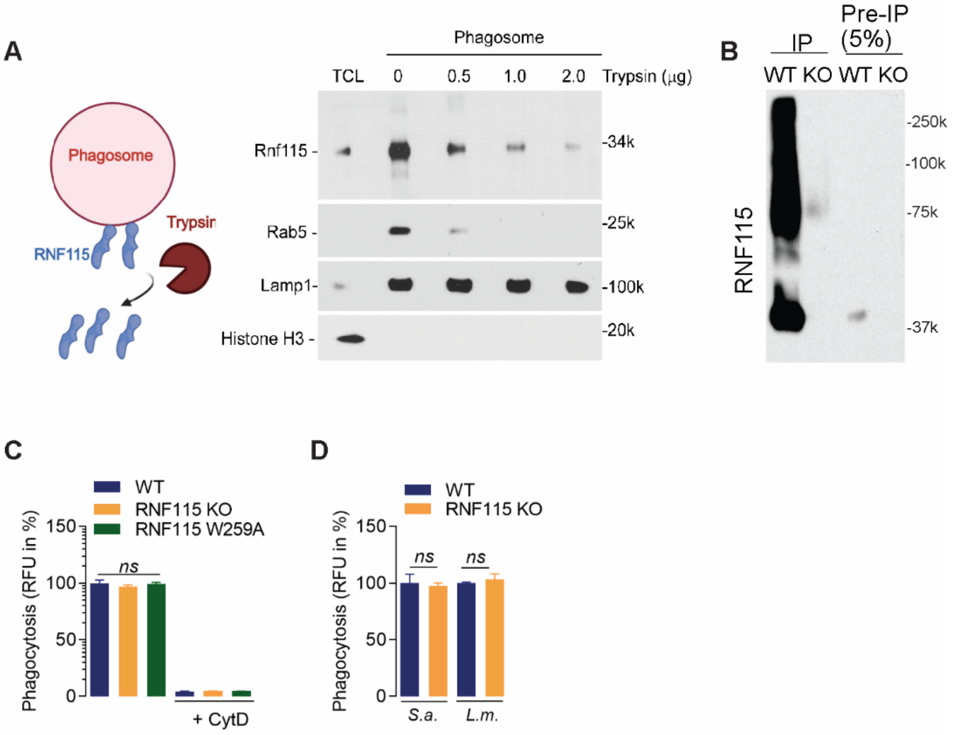


**Supplementary Figure 3: Phagosomal RNF115.** (A) RNF115 is located on the cytoplasmic side of phagosomes. Treatment of isolated phagosomes with increasing amounts of Trypsin shoes a reduction of RNF115. Rab5, a cytoplasmic membrane bound protein and LAMP1, a transmembrane protein, serve as controls. Histone H3 serves as purity control. Representative blot of two replicates. (B) Immunoprecipitation (IP) of RNF115 from WT and RNF115 knock-out RAW263.7 cells (KO) shows that RNF115 is absent in the KO cells. (C) Loss of RNF115 (KO) or mutation of the E2 binding site (W259A) does not affect uptake of carboxylated beads or (D) bacteria. Cytochalasin D (CytD) which inhibits phagocytosis, is used a negative control. RFU = relative fluorescence units. S.a. *Staphylococcus aureus*; L.m. *Listeria monocytogenes*.

#### Supplementary Figure 4

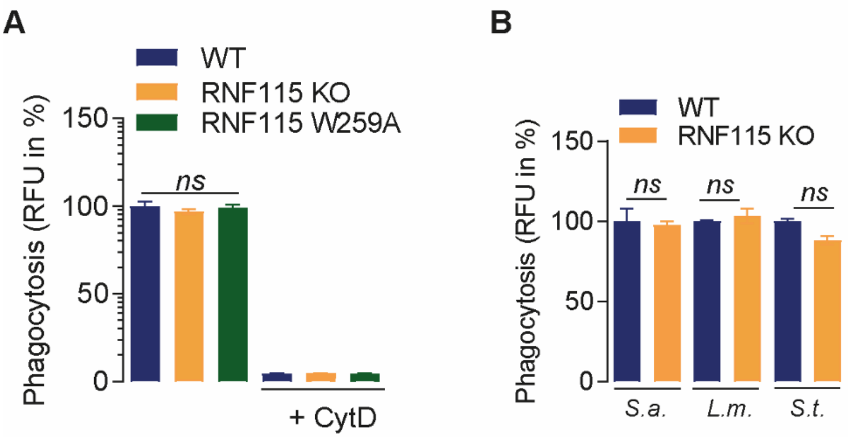


**Supplementary Figure 4: Loss of RNF115 does not affect phagocytosis** (A) Phagocytosis of 1 µm Silica beads is not affected by loss of RNF115 or expression of the “ligase-dead” RNF115 W259A in the KO strain. Cytochalasin D (CytD) serves as negative control as it blocks phagocytosis. (B) Phagocytosis of *Staphylococcus aureus* (*S.a.*), *Listeria monocytogenes* (*L.m.*) and *Salmonella* Typhimurium (*S.t.*) is not affected by loss of RNF115.
